# Pollen production, pollen viability and autofertility in faba bean (*Vicia faba* L.) and their relationship with realized paternal success

**DOI:** 10.1101/2023.11.28.568962

**Authors:** Lisa Brünjes, Wolfgang Link

**Affiliations:** Plant Breeding Methodology, Department of Crop Sciences, Georg-August Universität Göttingen, Carl-Sprengel-Weg 1, 37075 Göttingen, Germany, (LB), (WL)

**Keywords:** Faba bean, breeding, heterosis, pollen production, pollen viability, autofertility, paternal success, male reproductive success, impedance flow cytometry

## Abstract

In animal-pollinated plants, pollen dispersal depends on several plant and animal characteristics which may influence a plant’s paternal success. Different paternal success influences the genetic contribution of a genotype to the next generation. In breeding of partially allogamous faba bean (*Vicia faba* L.), synthetic populations are developed where equal contributions of genotypes to the next generation are desired to reduce inbreeding. Since direct assessments of paternity are elaborate and costly, we studied whether components of plant fitness such as pollen production and pollen viability can be used as estimates for paternity. In a field experiment and a caged outdoor pot experiment, a total of 18 genotypes (14 inbred lines, 4 F1 hybrids) of faba bean were evaluated for pollen production, pollen viability and autofertility. Pollen production was higher at the lower than at the upper inflorescences and we found mid-parent heterosis for this trait. The relative pollen viability was high (93 % – 97 % in pots, 88 % – 95 % in field) indicating that fertilization success is rather not limited by a low pollen quality. Only in the field, pollen of F1 hybrids was more viable than pollen of inbred lines. Autofertility ranged from 0 % – 98 %, with very marked average mid-parent heterosis for this trait. Autofertility did not seem to be related to either pollen production, pollen viability or paternal success. However, pollen production and pollen viability were highly correlated with paternal success. Hence, data on pollen production and viability might be useful in breeding of synthetic populations to choose parents with small differences in paternal successes, to reduce inbreeding and better exploit heterosis.

## 1. Introduction

The grain legume faba bean (*Vicia faba* L.) is a partially allogamous crop, showing an intermediate mode of reproduction between complete self-fertilization and complete cross-fertilization. In this respect, faba bean differs from most pulse crops of economic importance such as pea, soy or lentil, which are autogamous species. Several less commercially important pulses show a certain level of outcrossing, such as pigeon pea (*Cajanus cajan*) with 25-30 % (Saxena et al. 2021), white lupine (*Lupinus albus* L.) with 5-10 % (Faluyi and Williams 1981, Huyghe 1997), blue lupine (*Lupinus angustifolius* L.) with 0-12 % (Forbes et al. 1971), and yellow lupine (*Lupinus luteus*) with up to 40 % (Wallace et al 1954). In faba bean, however, the outcrossing rate can be 66 % or even higher (Gasim et al. 2004).

In faba bean, cross-fertilization fully depends on pollen transfer by pollinators, such as honey bees, bumble bees and solitary bees. In contrast, autogamous self-fertilization can be realized in two ways: First, as spontaneous selfing which requires a certain level of autofertility (see below) in the plant, and, second, as induced selfing, where a pollinator mechanically stimulates the flower (‘tripping’) thereby inducing the rupture of the stigma-covering membrane and thus pollen germination. Thus, the degree of cross-fertilization strongly varies affected by genetic factors, such as the general genotype, its autofertility and its level of inbreeding (Link 1990; Link and Ederer 1993; Gasim et al. 2004; Brünjes and Link 2021), and by environmental factors such as abundance, species composition and behavior of pollinators (Suso et al. 2001; Marzinzig et al. 2018) or geographical region (Suso and Moreno 1999).

To deal with this mode of reproduction in terms of a breeding strategy, synthetic cultivars (synthetics) are developed, which exploit both, the high performance of superior inbred lines and a certain share of heterosis due to the high level of outcrossing. In faba bean, synthetic cultivars are developed by selecting a limited number of inbred lines as parental components that are grown as mixed stand in spatial isolation, producing seed by open pollination. This generation is followed by several generations of seed multiplication under natural pollination without further selection, and the resulting seed is then sold to farmers (Brünjes and Link 2021). In such synthetics, the plants are more or less heterozygous and express a certain share of heterosis that will permanently stay in the synthetic population. Synthetic cultivars are genetically heterogeneous which supports yield stability and resistance to biotic and abiotic stresses (Link et al. 1994b; Stelling et al. 1994).

To minimize inbreeding and maximize the share of heterosis in synthetic populations, an equal contribution of parental components to the synthetic would be favorable. However, the success as pollen donors within the cross-fertilized seeds may differ between parental genotypes, as we have shown in a recent study (Brünjes and Link 2021). Differences in paternal outcrossing success lead to a higher level of inbreeding in synthetic populations than expected and thus a lower yield performance (for a discussion on this, see Brünjes and Link 2021). It would be favorable to combine those lines as parental components that do not differ in paternal outcrossing success. Then, inbreeding would be minimized and the exploitation of heterosis would be improved. However, directly assessing paternity is elaborate and costly, thus, characterizing a genotype for its paternal outcrossing success is unfeasible in practical breeding. Furthermore, is not known which plant characteristics contribute to the observed variation in paternal outcrossing success.

Paternal outcrossing success is difficult to assess. First, it requires large-scale paternity testing of many seeds (e.g. by SNP markers) comprising a high number of manually harvested seed, lab cost for marker analysis and a work-flow to interpret the data. Second, it is specific for a given set of genotypes, thus, the trait expression of a genotype not only depends on location and year but also on the other genotypes included in the set. Thus, it is unfeasible for breeders to assess this trait in their breeding material. Therefore, we might look for auxiliary traits, i.e. other reproductive traits that are possibly interrelated with paternal outcrossing success.

For instance, those genotypes which produce a higher number of pollen grains per flower might realize higher success rates as pollen donors to cross-fertilized seeds. In faba bean, pollen number per flower has been estimated before with reported results that varied strongly (Kambal et al.1976; Carré et al. 1994; Suso et al. 2008; Chen 2009; Bailes 2018; Aguilar-Benitez et al. 2022). A four-fold variation in pollen production has been found between 30 faba bean inbred lines by Bailes et al. (2018) and it has been shown that F1 hybrids produced more pollen grains per flower than their parental lines (Kambal et al. 1976; Chen 2009). The differential pollen production might not only be due to differences in plant material or the method of pollen counting, but also depend on the node from which the flowers were taken, which was not specified in these previous studies (see Carré et al. 1994 as an exception). Furthermore, no study on pollen production has been conducted on winter beans, yet.

Besides pollen production, different viability of the available pollen grains might also affect paternal outcrossing success. In the literature, the term pollen viability has been used to describe different aspects of pollen quality, such as stainability with certain dyes, germinability, pollen vigor or fertilization ability (for an overview, see Dafni and Firmage 2000). Pollen that is classified as viable not necessarily has the ability to germinate (e.g. due to unproper conditions) or fertilize the ovule (e.g., because of some sort of incompatibility) (Impe et al. 2020). Thus, we use viability as an operational term. While germinability is mostly assessed in vitro on plant-species specific culture media, mere viability is generally characterized by fluorescein diacetate (FDA) staining (Heslop-Harrison and Heslop-Harrison 1970) or other staining methods, such as acetocarmine or Alexander staining (Singh 2017), characterizing an intact cell membrane and denoting the presence of enzyme activity (Heslop-Harrison 1992). A more recent method to assess pollen viability is by impedance flow cytometry (IFC). The technology is based on a microfluidic chip, in which cells flow through and are exposed to an alternating electric current field in the radiofrequency range (0.1 – 30 MHz). The resulting impedance signals of the single cells are then used to characterize cell size, membrane integrity and cytoplasmic properties (Heidmann et al. 2016). High correlations were found between pollen-viability results of classical FDA staining and IFC in cucumber, tomato and sweet pepper (Heidmann et al. 2016), and in wheat (Bokshi et al. 2019; Impe et al. 2020). In faba bean, pollen viability was first assessed by Aguilar-Benitez et al. (2022) in two inbred lines using the acetocarmine staining.

Apart from pollen characteristics, paternal outcrossing success could be indirectly influenced by the ability of a plant to self-fertilize and set pods without any mechanical stimulation of the flower, termed autofertility (Drayner 1959). Autofertility in faba bean can range from 0 % to 100 % and has a clear heterotic component with inbred plants showing a lower autofertility than hybrid plants (Drayner 1959; Stoddard 1986; Link et al. 1994a). Previous studies used different parameters to describe autofertility, e.g. flowers per node, pods per node, pods per flower or seeds per pod, sometimes using all flowers of a plant (Lord and Heslop-Harrison 1984; Torres et al. 1993; Aguilar-Benitez et al. 2022), sometimes standardizing the number of flowers (Kambal et al. 1976; Link 1990; Chen 2009; Gasim et al. 2011; Puspitasari 2017) to account for the highly variable flower number in the faba bean species (50 to 300 per plant, Rowlands 1960).

In thus study, we aimed at auxiliary traits that can be easier assessed than paternal outcrossing success and might be used instead to improve faba bean breeding. We assessed pollen production, pollen viability and autofertility in faba bean to quantify the variation of these traits across a large set of genotypes and to identify whether mid-parent heterosis is present in these traits. Further, we studied how these traits are correlated to paternal outcrossing success and whether pollen production is influenced by the position of the sampled inflorescence. To this aim, we studied 18 genotypes of winter faba bean in two environments: first, a field experiment with eight genotypes where we assessed pollen production, pollen viability and paternal outcrossing success, and second, a caged outdoor pot-experiment with a larger set of 18 genotypes, where we assessed pollen production, pollen production across the flowering period of the plant, pollen viability and autofertility. To identify the impact of environment on pollen production and pollen viability, data obtained in the field and in the pot-experiment were related.

Under normal growing conditions, genotypes and plants would highly vary for the number of flowers and, if their fertilization was no restriction, for their maximum possible seed set (Rowlands 1960). One inflorescence bears three to eight flowers, but rarely produces more than four pods, and only about 25 % of fertilized flowers develop into pods (Stoddard 1986; Aufhammer and Götz-Lee 1991). This excess of flowers is an ecological strategy to attract pollinators and a common feature in mixed-mating species (Patrick and Stoddard 2010), however, it impedes an interpretation of the resulting number of pods per plant in terms of autofertility since even the most autofertile plant would not achieve 100 % pods per flower. In contrast, when only about one third of the flowers were left at the plant, all of them could develop into pods if autofertility was 100 % (Link 1990). As the available assimilates were distributed to a variable number of sinks, a differential flower number might further affect the number of pollen grains per flower or the pollen viability. Thus, when assessing autofertility in the pot-experiment, we decided to standardize flower number in order to eliminate potential differences caused by a different number of flowers per plant, and trimmed the plants (see Puspitasari 2017).

## 2. Material and Methods

### 2.1 Plant material

To assess pollen production and pollen viability, we grew eight faba bean genotypes in a field trial and the same eight genotypes plus ten additional genotypes in a caged outdoor pot-experiment. The plant material consisted of 12 winter faba bean inbred lines and four F1 hybrids (Table 1). The inbred lines were selected to show similar begin of flowering (max. difference two days) and to cover a wide range of autofertility levels (for further aspects, see see Brünjes and Link 2021). To create the F1 hybrids, parents with markedly different autofertility were selected. As reference checks for autofertility, we included the highly autofertile inbred line FAB172, which is of the *Vicia faba paucijuga* type, and the highly autosterile inbred line Diana07, which was bred from the outdated cultivar Diana. Genotypes 1 to 16 were winter faba beans and genotypes 17 and 18 were spring faba beans. The used winter bean genotypes set flowers and seed even if sown in spring, albeit some days later than spring types.

**Table 1.** Genotypes included in our study. Autofertility category “high” = at least seven pods or 15 seeds per untripped plant, “low” = max. one pod or two seeds per untripped plant, “medium” = in between low and high. Data on autofertility was taken from Puspitasari (2017) for genotypes 1 – 12, from Link (1990) for genotype 17, and from own unpublished field books for genotype 18. X = included in the corresponding experiment, − = not included

| Cont. number | Genotype | Autofertility category | Field experiment* | Caged outdoor pot-experiment |
| --- | --- | --- | --- | --- |
| 1 | S_003 | low | - | X |
| 2 | S_019 | medium | - | X |
| 3 | S_025 | high | - | X |
| 4 | S_035 | low | - | X |
| 5 | S_046 | low | X | X |
| 6 | S_085 | high | X | X |
| 7 | S_120 | low | - | X |
| 8 | S_145 | low | X | X |
| 9 | S_199 | low | X | X |
| 10 | S_217 | low | - | X |
| 11 | S_235 | low | X | X |
| 12 | Fam157 | medium | X | X |
| 13 | F1(S_019 x S_035) | medium x low | X | X |
| 14 | F1(S_025 x S_217) | high x low | X | X |
| 15 | F1(S_046 x S_085) | low x high | - | X |
| 16 | F1(S_199 x Fam157) | low x medium | - | X |
| 17 | Diana07 | low | - | X |
| 18 | VF172 | high | - | X |

\*Corresponds to set A in Brünjes and Link (2021)

### 2.2 Field experiment

To assess pollen production and viability under field conditions, we grew two F1 hybrids and six inbred lines (see Table 1) at a location West of Göttingen. These genotypes (referred to as “set A” in Brünjes and Link 2021) were a sub-set (“sub-set 1”) of the genotypes included in the caged outdoor pot-experiment. Embedded within a crop stand of faba bean, we had 12 rows composed of these eight genotypes with one single plant per genotype in each row. Each of these rows was thus treated as one replicate and genotypes were arranged randomly within each row. Plants were sown manually on March 15, 2016 in rows with 22 cm distance within the row and with alternating row-to-row distances (22 cm and 40 cm to better allow the access of persons). The plants in the field experiment were not standardized, e.g., neither tillers nor flowers were removed. When all single plants had started flowering, each tiller was labelled at that inflorescence which was then in full bloom (see Brünjes and Link 2021). On the next day, each single plant was sampled with one flower bud which was positioned one or two nodes above that labeled inflorescence; usually, this was the fourth or fifth inflorescence of the tiller. The samples were analyzed directly in the field for pollen production and pollen viability.

### 2.3 Caged outdoor pot-experiment

#### 2.3.1 Experimental design

Plants were grown in pots of 18 cm x 18 cm x 18 cm in bee-proof cabins that allowed a certain exposure of plants to rain and wind. To lessen the workload during flowering time and to balance the results against their dependency on actual temperature at flowering days, the the sowing dates were staggered. Thus, there were three sowing dates (February 3, February 12 and February 25, 2016), which would in field settings mean rather early sowing. Per sowing date, there were two replicates and two plants of each genotype per replicate. The two plants per replicate (i.e., the two pots) were directly neighbored and treated as one entry, while the entries in each replicate were randomized. Hence, across the three sowing dates, each genotype was included with a total of 12 plants. Plants were watered regularly and attentively, thereby preventing influence of water stress on pollen development.

#### 2.3.2 Plant standardization and bud sampling

The plants of the first sowing date were used to estimate autofertility, i.e. they were left untripped and displayed this trait via set of untripped pods. We kept only the two strongest tillers per plant. At the first eight inflorescences of tiller 1 and at the first four inflorescences of tiller 2, flower number was reduced to two, resulting in a total of 24 flowers per plant. All other tillers and inflorescences were carefully removed. Three nodes and leaves above the uppermost inflorescence, the tillers were chopped. Given this limited number of flowers and the abundance of water and nutrients, we assumed that plant assimilates sufficed to provide each flower with the option to develop into a pod if it was autofertile. If the number of seed-filled pods per plant was lower than the number of flowers per plant, this was hence taken as indication of a non-complete fertilization of these flowers. At harvest, the number of seeds and the number of pods of each plant was recorded. Then, the mean of the two neighboured single plants was calculated to obtain the mean values for each replicate, i.e., each entry.

The plants of sowing dates two and three were used to estimate pollen production and viability, i.e. flower buds were harvested and plants were not supposed to set pods. A standardization treatment similar to the one in sowing date 1 was applied to the plants, yet tillers 1 and 2 were both standardized to bear 10 inflorescences with four flowers each. Again, one entry was composed of two plants of the same genotype. To estimate pollen production (pollen grains produced per flower), we employed tiller 1 of the plants of all 18 genotypes in sowing dates 2 and 3. From each tiller, two buds from the lower (4^th^ and/or 5^th^) and two buds from the upper (7^th^ and/or 8^th^) inflorescences were collected (Table 2). To assess pollen viability, we used tiller 2 of two F1 hybrids and their four parental inbred lines in sowing date 3. Our aim was to sample each plant at inflorescences four to six, with a total of four samples per plant. However, due to very fast progression of blooming, final samples originated from inflorescences 5 to 13 (Table 2). To assess whether pollen production varies within a genotype across the flowering period, we used tiller 2 of the other two F1 hybrids that were not employed for analyzing pollen number, and their four parental inbred lines, in sowing date 3 (Table 2). At these tillers, each of the ten inflorescences was sampled with two buds. Buds were picked at the developmental stage that is optimal for emasculation, i.e. when the flower color turned from green to light white and anthers were shortly before dehiscing (Bond, Lawes and Poulsen 1980). To diminish possible effects of time of the day on bud development and pollen, buds in sowing dates 2 and 3 were picked at different hours per day.

**Table 2.** Traits evaluated in the caged outdoor pot-experiment. In each replicate, one entry is composed of two plants.

| Parameter | Autofertility | Pollen production | Pollen viability | Pollen production across inflorescences |
| --- | --- | --- | --- | --- |
| Sowing date | 1 | 2 and 3 | 3 | 3 |
| Number of genotypes | 18 | 18 | 6 (= sub-set 2) | 6 (= sub-set 3) |
| Genotypes | All | All | S_025,<br>S_046,<br>S_085,<br>S_217,<br>F1(S_025xS_217),<br>F1(S_046xS_085) | S_019,<br>S_035,<br>S_199,<br>Fam157,<br>F1(S_019xS_035),<br>F1(S_199xFam157) |
| Replicates | 2 | 4 | 2 | 2 |
| **Standardization** |  |  |  |  |
| Tillers | 2 | 2 | 2 | 2 |
| Inflorescences per tiller | 8 (tiller 1)<br>4 (tiller 2) | 10 | 10 | 10 |
| Flowers per inflorescence | 2 | 4 | 4 | 4 |
| **Sampling** |  |  |  |  |
| Employed tillers | None | Tiller 1 | Tiller 2 | Tiller 2 |
| Inflorescence position [1-10]* | None | 4 or 5 (LOW) and<br>7 or 8 (UP) | 5 to 10 | 1 to 10 |
| Buds sampled per tiller | None | 4 | 4 | 20 |

\* Lowest inflorescence = inflorescence position 1. Each node upwards was consecutively numbered with inflorescence position 2, 3, etc

### 2.4 Collection and preparation of pollen samples

Flower buds were harvested at one day prior to presumed anther dehiscing to ensure that all pollen grains of a flower were collected. Directly after bud picking, the keel (*carina*) was sliced open with tweezers and the correct stage of maturity was checked, i.e. anthers might not have dehisced to ensure that all pollen grains of a bud were gathered. All ten anthers of a flower were put into an Eppendorf tube and were either stored at −20 °C for later processing (test for pollen production) or analyzed directly within a few minutes after collection (test for pollen viability). It was assumed that counts of pollen production were not affected by freezing of samples, whereas freezing would have affected viability (Nath and Anderson 1975; Shivanna et al. 1991).

### 2.5 Impedance flow cytometry

Pollen production and pollen viability were estimated using impedance flow cytometry (IFC), the method was established for faba bean (pre-tests see Supplementary file 1). For each sample, the ten closed anthers of a flower bud were transferred into an Eppendorf tube, slightly squeezed to release the pollen grains and suspended into 2 ml of IFC buffer solution (AF 5, Amphasys, Lucerne, Switzerland). The suspension was filtered twice using 100 µm pore size and loaded to a 120 µm impedance chip of an impedance flow cytometer type Ampha Z30 (Amphasys, Lucerne, Switzerland). Pollen production and pollen viability were measured at 0.5 MHz employing the default settings (Trigger level 0.1 V, Modulation 3, Amplifier 6, Demodulation 2), while the pump was set to 300 rpm. The line gating was set to 330 phase angle and used as template in all samples. Measurements were analyzed using AmphaSoft version 1.2.4 (Amphasys, Lucerne, Switzerland). Data of samples with a concentration of at least 1,000 cells ml^-1^ and less than 15 % rejected cells of total cell counts entered the statistical analysis.

Estimates for pollen production (i.e., number of pollen grains per flower) were the number of accepted counts, whereas pollen viability was estimated as (i) the absolute number of viable pollen grains per flower, using the IFC-based counts of viable cells, and (ii) the percentage of viable cells in total number of IFC-accepted counts.

### 2.6 Statistical analysis

The statistical analysis was performed in R (version 4.1.2, R Core Team 2021) using the “lme4” package (Bates et al. 2015). Figures were produced employing the “ggplot2” package (Wickham 2016) and the “multcompView” package (Piepho 2004) to obtain compact letter displays. If not specified differently, significant differences are not presented based on a pre-defined threshold of an error margin, but as the resulting p-values themselves.

We checked the linear models for potential overdispersion of data and used residual plots, Q-Q plots and Shapiro-Wilk tests to validate the normal distribution of residuals. We performed Levene’s tests to validate the homogeneity of variance. Additionally, we performed residual diagnostics using the “DHARMa” package (Hartig 2022). The results of the model diagnostics are not reported here.

#### 2.6.1 Pollen production and pollen viability in the field experiment

In the field experiment, pollen production and pollen viability were assessed at two F1 hybrids and six inbred lines, sampled at inflorescence 4 or 5 in twelve replicates. Each genotype was tested with one sample (i.e., one flower bud) per replicate, thus, 96 samples were used in this experiment. Three reproductive traits were studied: pollen production (i.e., number of pollen grains per flower), pollen viability absolute (i.e., number of viable pollen grains per flower), and relative pollen viability (i.e., ratio of viable pollen grains to total pollen grains). We used linear models with genotypes and replicates as fixed factors to test for differences between genotypes in these traits. Tukey’s tests were performed to identify differences between genotypes in all three reproductive traits (p = 0.05).

#### 2.6.2 Pollen production in the caged outdoor pot-experiment

In the pot-experiment, pollen production was estimated by employing 18 genotypes in four replicates. Variation in pollen production between flowers of early or later inflorescences was accounted for by using two traits: the mean values of each entry for (i) the two inflorescences at the lower part of the tiller (trait “LOW”), and (ii) the two inflorescences at the upper part of the tiller (trait “UP”). The samples collected from the lower and upper part of four tillers of an entry were bulked accordingly. Pollen production LOW was estimated from an average number of 3.51 samples (0 to 7 samples) per entry and replicate. Pollen production UP was estimated with on average 3.04 samples (0 to 7 samples) per entry and replicate. A total of 472 samples were used in the analysis.

Of the 72 data points of this trait, there was one value missing for LOW (1.4 %) and nine missing values for UP (12.5 %). Estimates for the missing data were obtained by iterative calculations that minimize the residual mean square (Yates 1938; Healy and Westmacott 1956) using the software PLABSTAT (Version 3A, Utz 2011). The complete dataset including the substitute values was then used to calculate the mean values of traits LOW and UP, which was used as third trait, i.e. pollen production per flower of the total plant (trait “TOTAL”).

To estimate the effect of genotype on pollen production, we employed a linear mixed effects model with genotypes as fixed factor and replicates as random factor to explain the variance of number of pollen grains (either LOW, UP or TOTAL). The two sowing dates were not included as source of variation. We conducted analyses of variance and adjusted the residual degrees of freedom according to the number of missing values. To ensure a near-to normal distribution of residuals, we performed a boxcox transformation (package “MASS”, Venables and Ripley 2002), using one value for lambda for each of the three traits (LOW: lambda = 1.798, UP and TOTAL: lambda = 2). Figures show results from the original data while all statistical tests were done with the transformed data. In the transformed data, we identified two outliers for each trait pollen production LOW and pollen production UP (package “rstatix”, Kassambra 2021) which were kept in the dataset. Least significant differences were calculated with the adjusted residual degrees of freedom according to the number of missing values at an error margin of 0.05. To identify which inbred line differed significantly from any other inbred line, we performed Tukey’s HSD test. Welch’s t-test were employed to test the differences of mean values between lower and upper inflorescence positions.

To check whether there is mid-parent heterosis for pollen production at all, we performed paired t-tests and tested whether the mean of the F1 was greater than the mean of its corresponding two parental lines. Given that heterosis for pollen production was significantly present, we tested whether the four crosses differed in the size of that heterosis. To that aim, we performed analyses of variance for (i) the absolute heterosis (pollen number superiority of F1 over their parental inbred lines), estimated with the boxcox-transformed values, and (ii) the relative heterosis (relative superiority of the F1 to the mean of its parents, Melchinger et al. 1994), estimated with the original values. Subsequently, Tukey’s HSD tests were employed to identify which of the crosses showed more heterosis than another cross. Finally, to identify for each cross separately if that particular cross showed heterosis for pollen production, we tested in each single cross whether the F1 hybrid was greater in pollen production than the mean of its parents, using Welch’s t-test.

#### 2.6.3 Pollen viability in the caged outdoor pot-experiment

Pollen viability was assessed in the pot experiment at two F1 hybrids and their four parental inbred lines in one sowing date. We used 78 samples (flower buds) in this experiment; each genotype was tested with on average 6.5 samples (5 to 8 samples) per replicate. Along with data of viability, we obtained data of pollen production. The following traits were considered: pollen production, pollen viability absolute, and relative pollen viability. To test for differences between genotypes in these traits, generalized linear models (GLM) were performed with genotypes and replicates as fixed factors. For the traits “pollen production” and “pollen viability absolute”, GLMs were used assuming negative binomial errors with a log link function. For the trait “relative pollen viability”, binomial errors with a logit function were used and the response variable was defined as a matrix where the first column was the number of viable pollen grains and the second column was number of non-viable pollen grains. Here, quasi-binomial errors were used due to overdispersion of the data. Tukey’s tests were performed to identify differences between genotypes in all three reproductive traits (p = 0.05).

#### 2.6.4 Pollen production across inflorescences in the caged outdoor pot-experiment

For the estimation of pollen production across inflorescences, we employed two F1 hybrids and their four parental inbred in two replicates. The intention was to obtain four buds (=samples) per entry at each of the ten inflorescence positions, which were used to calculate ten mean values per inflorescence position (each value an average of four buds). However, in 62.5 % (75 of 120 inflorescence positions), pollen data of less than four buds per inflorescence position was used, either because samples were discarded during quality control (concentration less than 1,000 cells/ml or a rejection rate higher than 15 %) or because buds were sampled too late, hence anthers had already opened in such buds or because the plant did not produce any buds at the particular node.

To calculate mean values per inflorescence position, we dealt with discarded and with missed buds as follows: Differences between buds at the same inflorescence position (grown at either inflorescence of the two neighbored single plants) were assumed to be random. Thus, the result of one inflorescence position was the average of the analyzed buds, be it one, two, three or four. If zero buds were obtained from a inflorescence position, we employed the values of the other replicate (block) yet duly respecting the block effect. Twice, there weren’t any values for existing buds in any of the two replicates. There, we took the mean value of all other inflorescence positions of that genotype (i.e. of the two neighbored single plants) as an estimate for the missing inflorescence. In contrast to quality-discarded buds and to too-late-picked buds, we treated non-existing buds in a different way. Inflorescence positions with zero buds, i.e. with non-existing buds, occurred in 20.4 % of the 480 intended-to-be-picked buds. Here, the number of pollen grains was taken as zero. In 3.3 % of the data, data on all four buds of an inflorescence position was missing, hence we adjusted the degrees of freedom in the analysis. On average, 2.41 buds per inflorescence position entered the analysis.

The inflorescence positions were obviously not randomized at the plant. To nevertheless estimate the effect of inflorescence position on pollen production, we used a split-plot design (package “agricolae”, de Mendiburu and Yaseen 2020) and conducted analyses of variance with genotypes as the effect of the main plot treatment (fixed factor), replicates as the block effect (random factor), the genotype-replicate interaction as the main plot error, inflorescence position as the effect of the subplot treatment (fixed factor), the genotype × inflorescence-position interaction and the three-way interaction of genotypes, replicates and inflorescence position as the subplot error. We conducted analyses of variance and adjusted the residual degrees of freedom of the residuals according to the number of missing values. To identify differences in pollen production between inflorescence positions, we employed Tukey’s HSD tests (p = 0.05).

#### 2.6.5 Autofertility in the caged outdoor pot-experiment

As measure for autofertility, we took the rate of fertilization (Link 1990), calculated as the ratio of number of seed-containing pods to the total number of untripped flowers, which gave an indirect score of the proportion of flowers that self-fertilized without being tripped. This trait lies within the range of 0 % to 100 %. In this study, 100 % rate of fertilization equaled 24 flowers that developed into 24 pods. We supposed that the more-autosterile plants do not only have fewer pods than more-autofertile ones, but that they additionally had fewer seeds within these few pods. Therefore, the trait seeds per pod was used as complement to rate of fertilization. Seeds per pod was not taken as synonym of autofertility, but as additional trait to recognize that alongside a varying pod-set based autofertility, also the seed-filling of pods varied. The trait seeds per pod did not follow a fixed scale, although more than four seeds per pod occur very rarely. It was not assessed pod-wise, but estimated as the ratio of total number of seeds per plant to the total number of pods per plant. If an entry had no pod, seeds per pod was set to zero.

To analyze the differences between genotypes in both traits rate of fertilization and seeds per pod, generalized linear models (GLM) were employed with genotype and replicate as fixed factors. To estimate rate of fertilization, a GLM with a binomial distribution was calculated. Here, the response variable was a matrix where the first column was the number of pods and the second column was the number of flowers minus the number of pods. To analyze seeds per pod, a GLM with a normal distribution was employed.

Twelve inbred lines that were studied here (cont. number 1–12 in Table 1) had been studied in previous years 2013-2015 using the same experimental setup employing different standardization methods which had no effect on rate of fertilization (Puspitasari 2017). To estimate rate of fertilization across four years, a generalized linear mixed model was used with genotype and year as fixed factors and replicate nested within standardization method nested within year as random factor. Again, the response variable was a matrix of binomial type (see above). Thereby, we were able to compare our data on pollen-related traits of one year to more robust data on rate of fertilization across four years.

## 3. Results

### 3.1 Pollen production and pollen viability in the field experiment

In the field experiment, we studied six inbred lines and two F1 hybrids and estimated pollen production, absolute pollen viability and relative pollen viability directly in the field. Pollen production ranged from 12000 to 20200 pollen grains per flower with a mean of 15400 in inbred lines and 18400 in F1 hybrids. F1 hybrids were significantly higher in pollen production (Welch’s two-sample t-test, t = 4.87, df = 40.81, p-value < 0.001). The two F1 hybrids differed significantly in pollen production (Tukey test, p-value < 0.001), and we found a large variation in pollen production of the inbred lines along with several significant differences (Fig. 1a). The standard error of the least square means was 741 pollen grains.

**Fig. 1.**
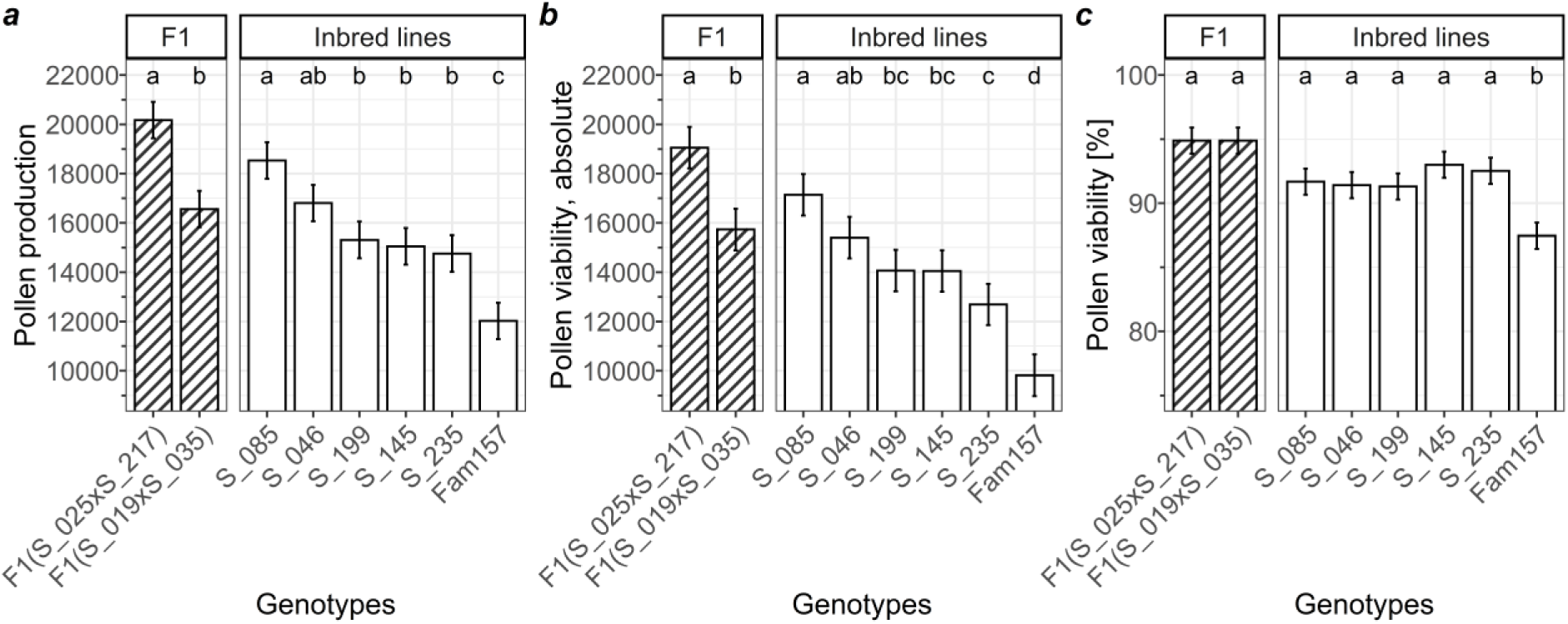
Reproductive traits of two F1 hybrids and six inbred lines grown in the field experiment: **a** Pollen production (pollen grains per flower), **b** Pollen viability (viable pollen grains per flower), and **c** Relative pollen viability (percentage of viable pollen grains per flower). Error bars show standard errors of least square means. Different letters show significant differences among least square means (p=0.05, Tukey’s HSD test).

The absolute number of viable pollen grains per flower bud ranged from 9800 to 19100 with a mean of 13900 in inbred lines and 17400 in F1 hybrids. F1 hybrids were significantly higher in absolute pollen viability than inbred lines (Welch’s two-sample t-test, t = 5.53, df = 45.04, p-value < 0.001). The two F1 hybrids differed significantly in absolute pollen viability (Tukey test, p-value < 0.001), and we found a large variation in absolute pollen viability of the inbred lines along with several significant differences (Fig. 1b). The standard error of the least square means was 841 viable pollen grains.

The percentage of viable pollen grains ranged from 87.5 % to 94.9 % with a mean of 89.9 % in inbred lines and 94.6 % in F1 hybrids. F1 hybrids were significantly higher in relative pollen viability than inbred lines (Welch’s two-sample t-test, t = 6.82, df = 51.37, p-value < 0.001). The two F1 hybrids did not differ in relative pollen viability (Tukey test, p-value = 1), and we found one of the inbred lines to be significantly different from all other lines (Fig. 1c). See Supplementary file 2, Supplementary Table 1 for all trait estimates per genotype.

In sum, the eight genotypes varied strongly and significantly in pollen production as well as in absolute and relative pollen viability. Although the F1 hybrids as a group had a higher pollen production and absolute pollen viability than the inbred lines, this was not the case for each single hybrid. One inbred line (Fam 157) was significantly lower than the other genotypes in all three traits. Genotype had a strong effect on pollen production and pollen viability in the field experiment, on an overall high level of these traits.

### 3.2 Pollen production in the caged outdoor pot-experiment

#### 3.2.1 Pollen production depending on the position of the flower at the plant

We estimated pollen production in 14 inbred lines and four F1 hybrids at inflorescence position 4 and 5 (LOW), at inflorescence position 7 and 8 (UP), and at the total plant (TOTAL), calculated as the mean of LOW and UP. In the group of all 18 genotypes, mean pollen production LOW (minimum, maximum) was 21800 (17900, 25800) pollen grains per flower. Pollen production UP and TOTAL were 20500 (16600 to 23900) and 21100 (17700 to 24900) pollen grains per flower, respectively (Supplementary file 2, Supplementary Table 2). The 14 inbred lines showed a large variation in pollen production (Fig. 2) with several significant differences (Tukey’s HSD test, with p=0.05). For example, for pollen production TOTAL, the inbred line S_019 differed significantly from lines S_199, S_085 and S_145. Further results on significance are not shown.

**Fig. 2.**
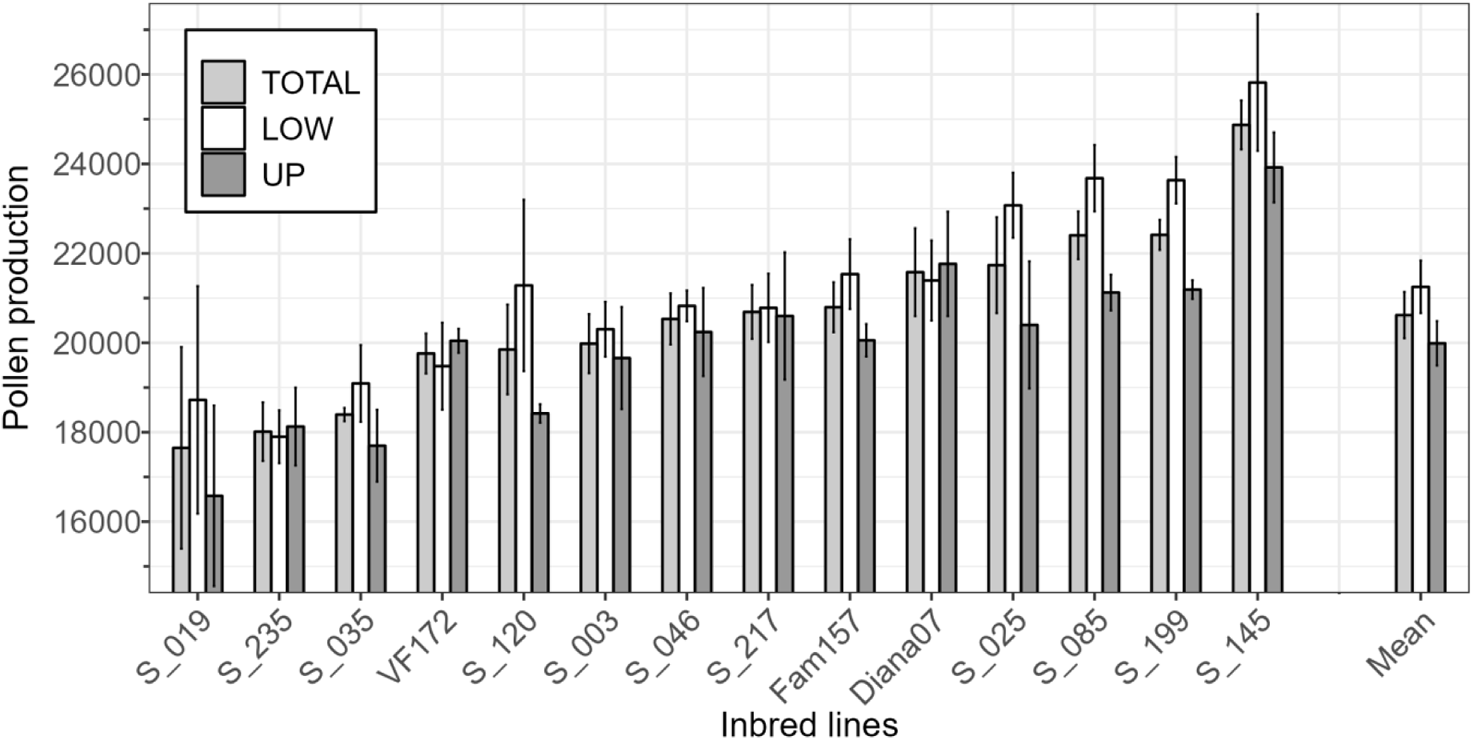
Pollen production (pollen grains per flower) of the 14 inbred lines as the average of the total plant (TOTAL), at the 4^th^ or 5^th^ inflorescence position (LOW) and at the 7^th^ or 8^th^ inflorescence position (UP). Note that the inbred lines are sorted by pollen production TOTAL, which is the mean of LOW and UP. Error bars show standard error of the mean, i.e., across the four replicates for the inbred lines and across the mean values of inbred lines for the overall means, respectively.

The inbred lines with the highest pollen production (e.g., S_145, S_085, S_199) were similar in pollen production to the F1 hybrids. In all three traits, inbred line S_145 showed the highest pollen production of all tested genotypes, i.e. even higher than any of the F1 hybrids (Supplementary file 2, Supplementary Table 2). The analyses of variance showed significant differences between genotypes for all traits LOW, UP and TOTAL (Supplementary file 2, Supplementary Tables 3, 4 and 5).

Generally, genotypes had higher pollen production LOW than UP or TOTAL but genotypes with high pollen production LOW did not always concomitantly have high pollen production UP. Therefore, we tested the relationship between traits LOW, UP and TOTAL and found a high significant correlation between pollen production LOW and UP (Fig. 3a), both calculated across all 18 genotypes (r = 0.82, p-value < 0.001) and across only the inbred lines (r = 0.82, p-value <0.001), as well as between pollen production LOW and TOTAL (Fig. 3b) calculated across all 18 genotypes (r = 0.96, p-value < 0.001) and across only the inbred lines (r = 0.97, p-value <0.001). Thus, the heterotic group-effect involved in the correlation was marginal. Overall, pollen production LOW was strongly and positively related to pollen production UP and TOTAL.

**Fig. 3.**
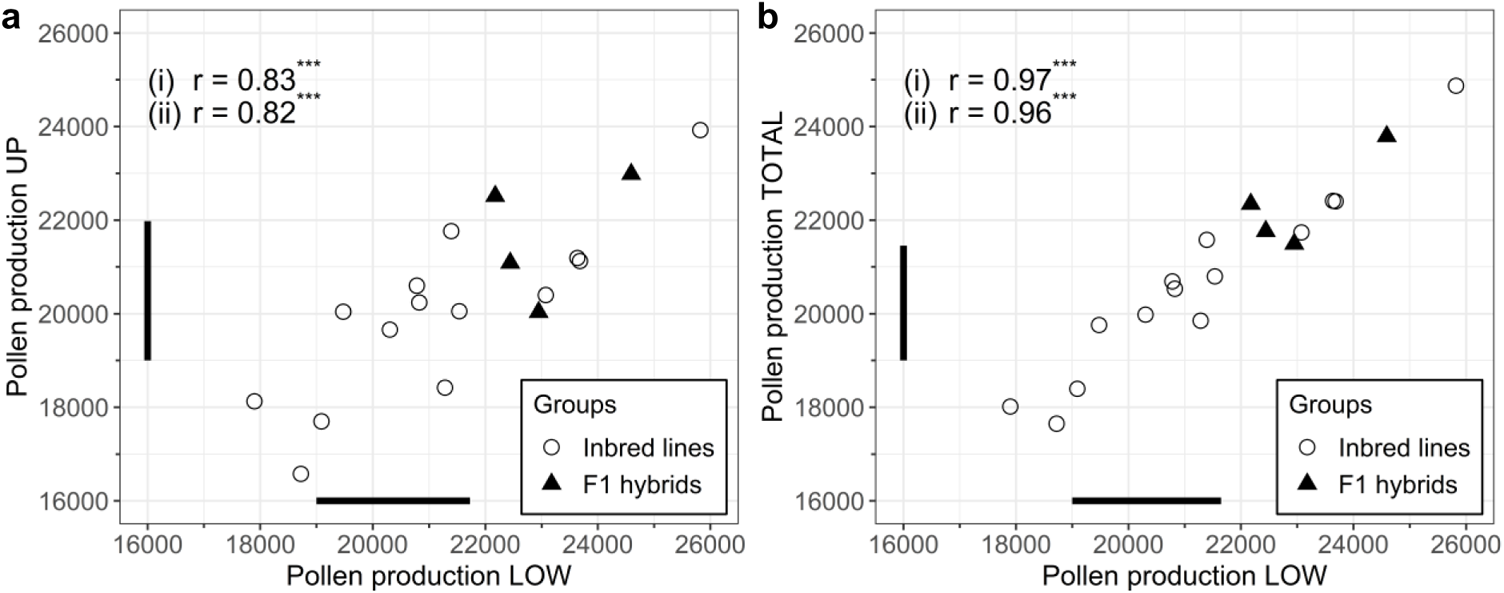
Correlation of pollen production LOW with **a** pollen production UP, and **b** pollen production TOTAL. Correlation coefficients were calculated based on Pearson’s product moment correlation coefficient with the boxcox-transformed data for (i) only inbred lines, (ii) all genotypes. Horizontal and vertical bars show the size of the least significant difference (with p=0.05) for **a** all 18 genotypes, and **b** only the 14 inbred lines (*** = significant at p < 0.001)

Does pollen production depend on the position of the sampled inflorescence at the plant and does pollen production LOW differ from pollen production UP? To answer that question, we have to shift to another perspective, no longer considering LOW and UP as two different traits, but considering LOW and UP as two levels of a treatment (“lower inflorescence position” and “upper inflorescence position”). The 18 genotypes showed a higher pollen production at the lower inflorescence position than at the upper inflorescence position (Welch’s t-test, t=-11.77, d.f.=8.27, p-value <0.001, performed with the mean values across the 4 replicates). Thus, across all 18 genotypes, pollen production was higher at the lower than at the upper inflorescence position.

To test if the effect of inflorescence position on pollen production was different in F1 hybrids than in inbred lines, we used data of only the four F1 hybrids and the eight parental inbred lines. Pollen production differed significantly between the lower and the upper inflorescence positions in the overall mean of F1 hybrids (Welch’s t-test, t = −13.31, df = 3.07, p-value <0.001) and in the overall mean of the eight inbred lines (Welch’s t-test, t = −14.95, df = 7.43, p-value < 0.001), see Fig. 4. It could therefore be concluded that indeed, pollen production depended on the position of the flower at the plant, with a higher pollen production at the lower than at the upper inflorescence positions. The effect of inflorescence position on pollen production was similar in F1 hybrids and inbred lines.

**Fig. 4.**
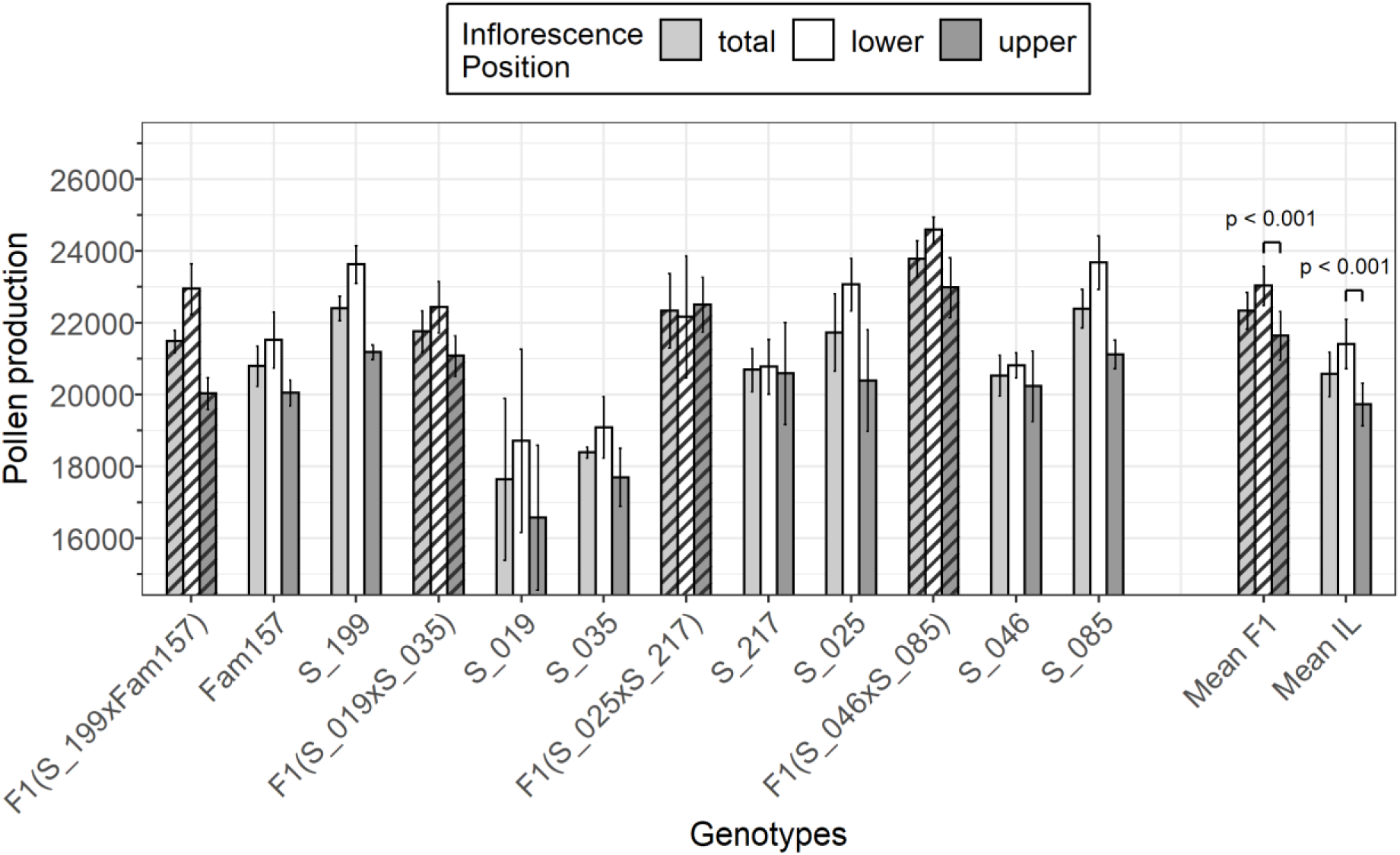
Pollen production (pollen grains per flower) of the four F1 hybrids (striped) alongside their parental inbred lines (not striped) at the total plant, at the lower inflorescence positions and at the upper inflorescence positions. Note that the four F1 hybrids are sorted by pollen production of the total plant, which is the mean of the values for pollen production at the lower and upper inflorescence positions. Error bars show standard error of the mean. For the genotypes, standard errors were calculated across the four replicates; for means of four F1 and mean of eight inbred lines (on the right), standard errors were calculated across the overall means of the genotypes of the respective group. The differences of mean values between lower and upper inflorescence position were tested with Welch’s t-test

#### 3.2.2 Heterosis for pollen production

As pollen production seems to be larger in F1 hybrids than in parental inbred inbred lines (see overall means in Fig. 4), we tested the four crosses (cont. number 13–16 in Table 1) for mid-parent heterosis for pollen production. Significant absolute heterosis was found for pollen production LOW (t-test, t = 2.55, df = 15, p-value = 0.011), UP (t = 3.22, df = 15, p-value = 0.003) and TOTAL (t = 3.06, df = 15, p-value = 0.004). The heterosis was larger for pollen production UP than for pollen production LOW (absolute heterosis: 1917 ± 938 (pollen grains per flower ± SE) vs. 1621 ± 799, respectively; relative heterosis: 0.107 ± 0.055 vs. 0.086 ± 0.046, respectively). However, the values varied a lot depending on the specific cross and replicate (Table 3).

**Table 3.**
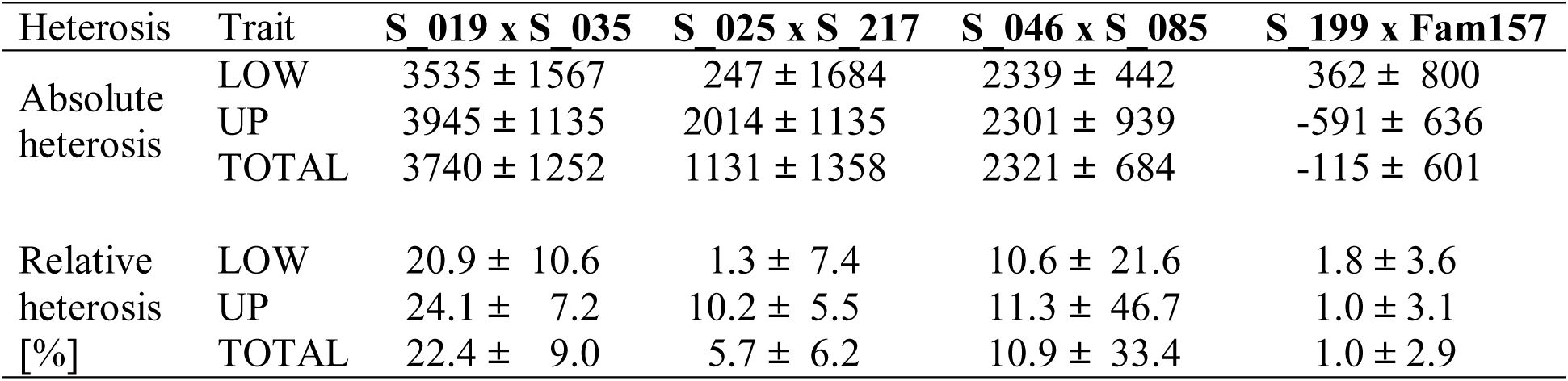
Heterosis for pollen production LOW, UP and TOTAL in the four crosses, in absolute heterosis (pollen number superiority) and in relative heterosis (%). Data are mean values and standard errors of the mean

| Heterosis | Trait | S_019 x S_035 | S_025 x S_217 | S_046 x S_085 | S_199 x Fam157 |
| --- | --- | --- | --- | --- | --- |
| Absolute heterosis | LOW | $3535 \pm 1567$ | $247 \pm 1684$ | $2339 \pm 442$ | $362 \pm 800$ |
| | UP | $3945 \pm 1135$ | $2014 \pm 1135$ | $2301 \pm 939$ | $-591 \pm 636$ |
| | TOTAL | $3740 \pm 1252$ | $1131 \pm 1358$ | $2321 \pm 684$ | $-115 \pm 601$ |
| Relative heterosis [%] | LOW | $20.9 \pm 10.6$ | $1.3 \pm 7.4$ | $10.6 \pm 21.6$ | $1.8 \pm 3.6$ |
| | UP | $24.1 \pm 7.2$ | $10.2 \pm 5.5$ | $11.3 \pm 46.7$ | $1.0 \pm 3.1$ |
| | TOTAL | $22.4 \pm 9.0$ | $5.7 \pm 6.2$ | $10.9 \pm 33.4$ | $1.0 \pm 2.9$ |

Given the existence of heterosis for pollen production, we further examined whether the four crosses as a group differed in size of heterosis. The crosses differed in size of heterosis for pollen production UP and TOTAL, but not so in pollen production LOW, both in absolute and relative heterosis (Analyses of variance, Supplementary file 2, Supplementary Tables 6 to 11). Finally, when testing for each cross separately whether that specific cross had significant mid-parent heterosis for pollen production, absolute heterosis was found for pollen production LOW, UP or TOTAL in the crosses S_019 x S_035 and S_046 x S_085 (Welch’s t-test, p<0.05; Supplementary file 2, Supplementary Table 12), but not so in the crosses S_025 x S_0217 and S_199 x Fam157. Thus, it could be concluded that mid-parent heterosis for pollen production existed in the tested four crosses while its size depended on the cross and on the position of the sampled inflorescences.

### 3.3 Pollen viability in the caged outdoor pot-experiment

Pollen viability was assessed at two F1 hybrids and their four parental inbred lines in one sowing date. Alongside viability, data on pollen production was obtained which was used as additional comparison between experiments. In the viability experiment, pollen production ranged from 8500 to 16200 pollen grains per flower, which was lower than pollen production of the same genotypes tested in the previous experiment (see section 3.2.1 and Supplementary file 2, Supplementary Table 2). The absolute number of viable pollen grains per flower was strongly correlated to pollen production (r = 1, p-value < 0.001) and only slightly lower than pollen production, as the relative pollen viability was high, ranging from 92.8 % to 96.6 % (Table 4). The position of the sampled inflorescences was on average 9.4 and thus farther up the tiller than in the experiment on pollen production (section 3.2.1). The position of sampled inflorescences did not differ significantly between the genotypes (Tukey’s HSD test, p = 0.05, Table 4).

**Table 4.**
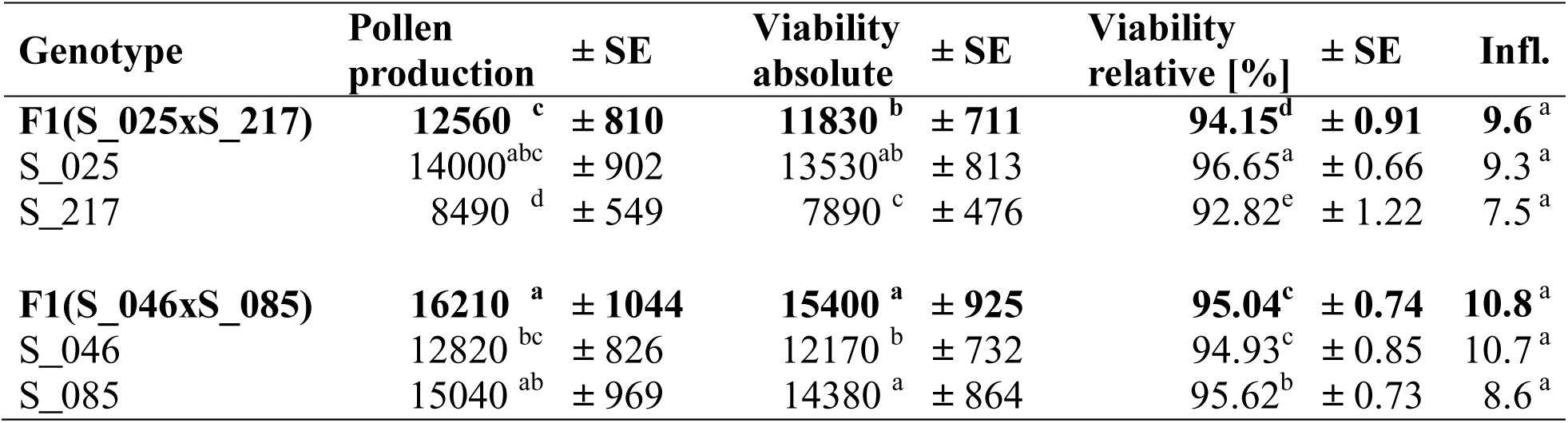
Pollen production, pollen viability absolute and viability relative of six genotypes in the caged outdoor pot-experiment. SE = Standard error of the mean, Infl. = mean position of sampled inflorescences. Different letters show significant differences among least square means (p=0.05, Tukey’s HSD test)

We found significant differences in absolute viability between all genotypes (Tukey’s HSD test, p=0.05, Table 4), as well as within the group of the two F1 and within the group of inbred lines (results not shown). However, the group of F1 hybrids did not differ in from the group of inbred lines (Welch’s t-test, p-value = 0.29). Relative pollen viability [%] significantly differed among all genotypes (Tukey’s HSD test, p=0.05, Table 4), within the group of two F1 hybrids (Tukey’s HSD test, p-value= 0.009), and within the group of inbred lines (Tukey’s HSD test, all four inbred lines were different from each other at p = 0.05). Again, the group of F1 hybrids did not differ in from the group of inbred lines (Welch’s t-test, p-value = 0.51).

It can be resumed that, as in the field experiment, relative pollen viability of the tested genotypes was very high. Hence, absolute pollen viability and pollen production were strongly and positively correlated. The genotypes differed significantly in relative pollen viability and the F1 hybrids were not necessarily higher in pollen viability than the inbred lines.

### 3.4 Pollen production across inflorescences in the caged outdoor pot-experiment

To study pollen production across the range of inflorescence positions of a tiller, we assessed the pollen grain numbers per flower at ten inflorescences of two F1 hybrids and their four parental inbred lines. The analysis of variance showed a significant main effect of the ten inflorescences across genotypes. Further, significant interactions of genotypes with replicates were found, as well as interactions of inflorescences with genotypes. There was no significant effect of genotypes (Table 5).

**Table 5.** Analysis of variance of pollen production per inflorescence, evaluated as a split plot design.

| Source of variance | d.f. | Mean Squares | F-value | p-value |
| --- | --- | --- | --- | --- |
| Genotype (G) | 5 | 1113 *10<sup>5</sup> | 1.63 | 0.303 |
| Replicate (R) | 1 | 1067 *10<sup>5</sup> | 1.56 | 0.267 |
| GxR | 5 | 684 *10<sup>5</sup> | 5.62 | < 0.001 |
| Inflorescence (I) | 9 | 3260 *10<sup>5</sup> | 26.79 | < 0.001 |
| IxG | 44 | 490 *10<sup>5</sup> | 4.12 | < 0.001 |
| Error | 51 | 115 *10<sup>5</sup> |  |  |

While the mean pollen production across all six genotypes was over 20000 pollen grains per flower at inflorescences 1 to 4, it decreased continuously to about 7000 pollen grains at inflorescence 10 (Fig. 5). Pollen production per inflorescence for each of the six genotypes is shown in Supplementary file 2, Supplementary Fig. 1 to 6.

**Fig. 5.**
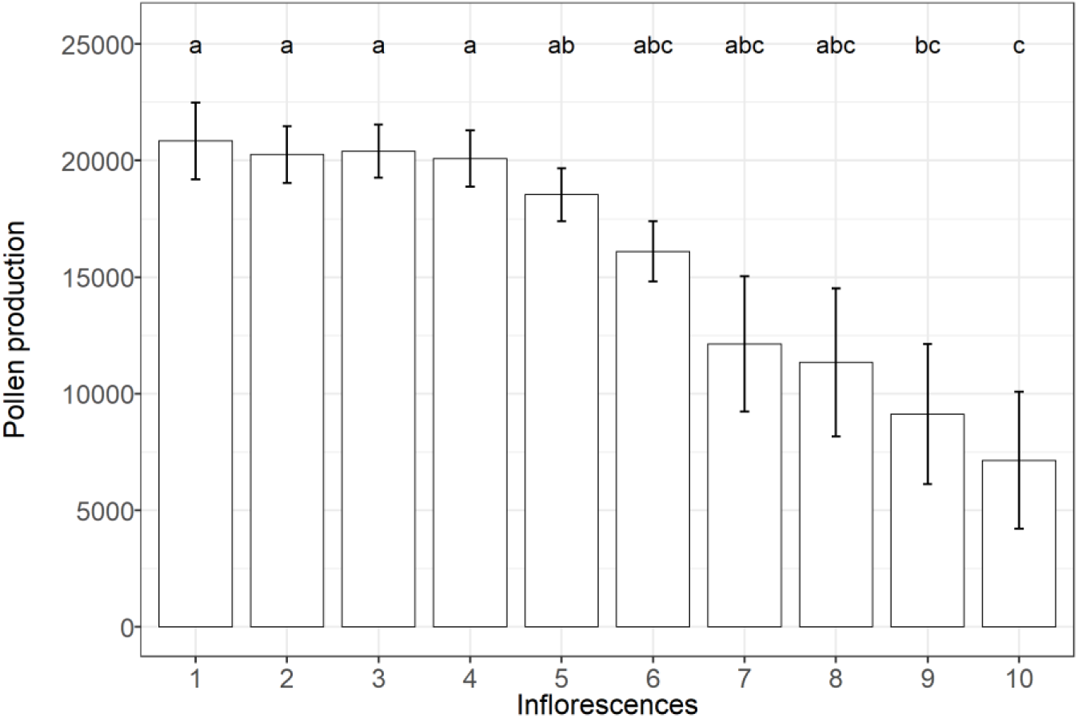
Pollen production (pollen grains per flower) at inflorescence positions 1 to 10. Shown are the mean values estimated at two F1 hybrids and their four parental inbred lines. Error bars show standard error of the mean. Different letters show significant differences among means (p=0.05, Tukey’s HSD test)

Hence, it can be concluded that inflorescence position had an effect on pollen production. More pollen grains per flower were found at the inflorescences at the bottom of a tiller than at the inflorescences further up the tiller.

### 3.5 Autofertility

We estimated autofertility of 18 genotypes as the rate of fertilization, i.e. as proportion of flowers that self-fertilized without being tripped. In our material of inbred lines and F1 hybrids, this trait ranged from 0 % to 97.9 % (Fig. 6; Supplementary file 2, Supplementary Table 13). The autofertility categories of Table 1 were mostly matched by the findings with our experiment. The three inbred lines previously classified as “highly autofertile” (S_025, S_085, VF172) were indeed high in rate of fertilization. S_019 and Fam157 were indeed “medium”, and all other inbred lines were lower. One deviation was line S_120, which was classified as “low”, but was one of the four inbred lines with the highest autofertility in our experiment.

**Fig. 6.**
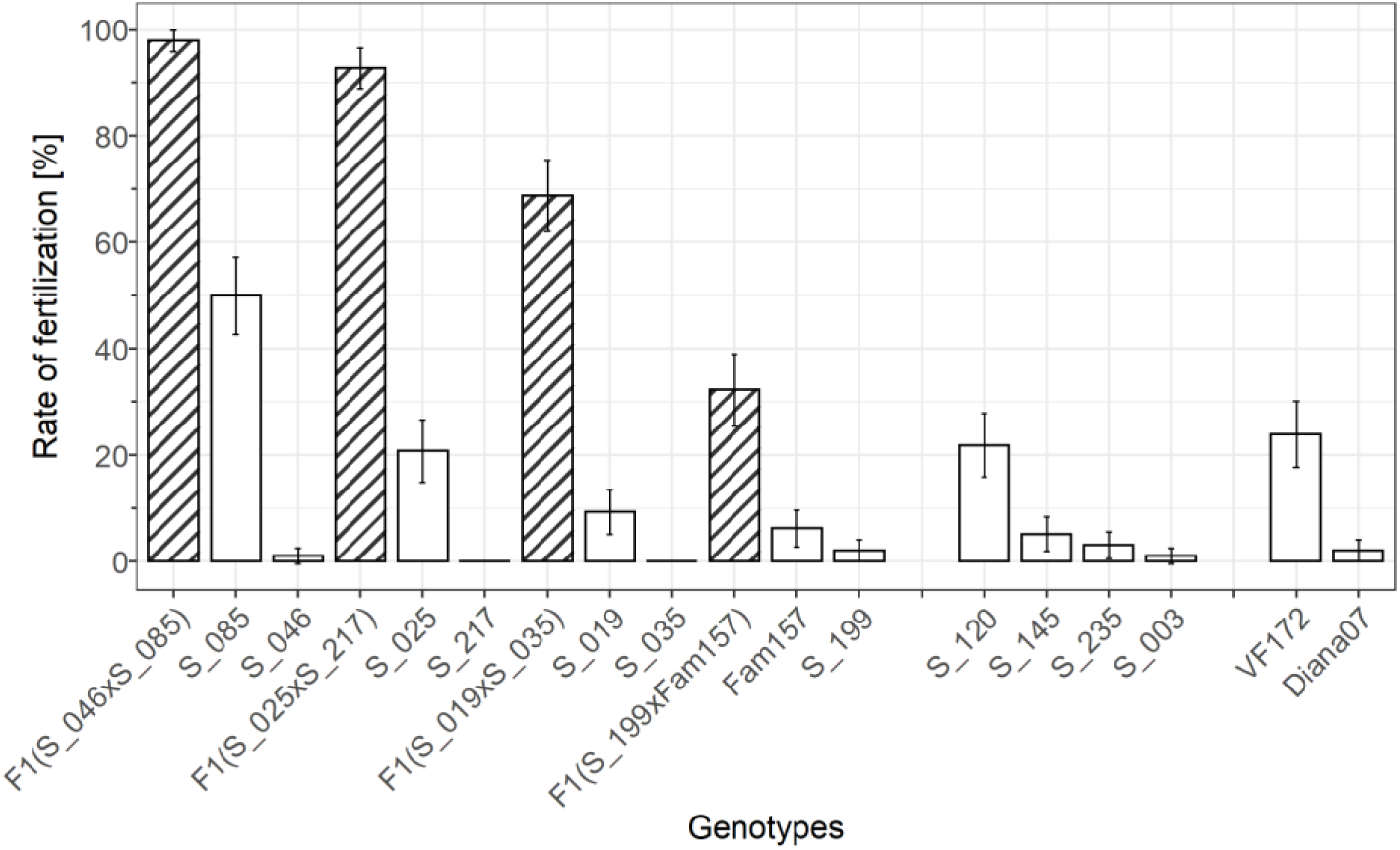
Rate of fertilization [%] of the four F1 hybrids and their parental lines, of four additional lines, and of two checks (high autofertility: VF172, low autofertility: Diana07). Note that the F1 hybrids were created by crossing two lines with diverging autofertility. Error bars show standard error of the mean

The F1 hybrids were created by crossing two lines differing in their autofertility. F1 hybrids had a higher rate of fertilization than any inbred line (mean rate of fertilization in F1 hybrids: 72.92 %, mean in all 14 inbred lines: 10.49 %). The higher autofertility in F1 hybrids was statistically significant (Welch t-test, t=6.16, df=8.04, p-value < 0.001). Within each of the two groups, Tukey HSD tests showed significant differences between genotypes in their rate of fertilization. Within the four F1 hybrids, all genotypes were significantly different from each other (Tukey HSD test, p < 0.001). Within the group of 14 inbred lines, most genotypes were significantly different from each other, except S_035 and S_217, S_003 and S_046, and S_199 and Diana07 (Tukey HSD test, with p=0.05).

Hence, the tested genotypes varied strongly in autofertility. Within both groups, inbred lines and F1 hybrids, genotypes strongly varied in autofertility. The F1 hybrids generally were autofertile (32 % to 98 %) whereas some inbred lines were autosterile and others were autofertile (0 % to 50 %). Thus, our tested material showed high variation for autofertility.

As autofertile plants might not only bear more pods than more-autosterile plants, but also have more seeds within the larger number of pods, we further analyzed seeds per pod as a complementary trait to autofertility. F1 hybrids bore more seeds per pod than inbred lines (mean in F1 hybrids: 3.04, mean in all 14 inbred lines: 1.51; Fig. 7), which was a significant difference (Welch t-test, t=6.25, df=22.12, p-value < 0.001). Within the group of F1 hybrids, there was no significant difference between genotypes in number of seeds per pod (Tukey HSD test, with p=0.05). Within the group of inbred lines, the two autosterile lines S_035 and S_217 were different from all other inbred lines except S_046, S_199 and Diana07. The four lines with the highest number of seeds per pod (S_025, S_120, S_235, Fam157) were different from the five lines with the lowest number of seeds per pod (S_035, S_046, S_199, S_217, Diana07. Tukey HSD test, with p=0.05).

**Fig. 7.**
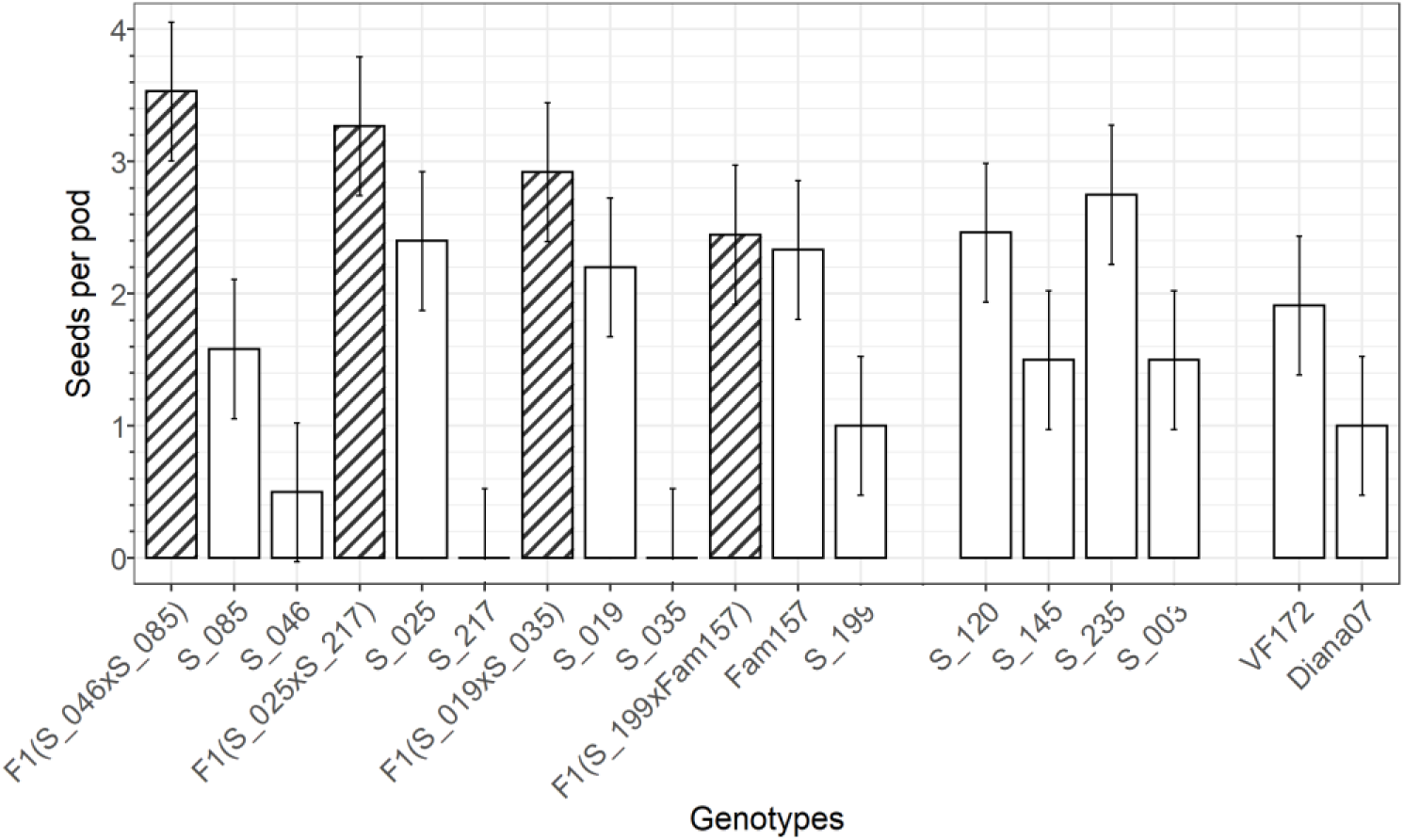
Seeds per pod of the four F1 hybrids and their parental lines, of four additional lines, and of two checks (high autofertility: VF172, low autofertility: Diana07). Error bars show standard error of the mean

Given the differences between F1 hybrids and the 14 inbred lines in rate of fertilization and seeds per pod, we checked whether there was heterosis for both traits. We used paired t-tests and tested for all four crosses whether the mean of the F1 was greater than the mean of its two parental lines, in each replicate (n=8). We found that hybrids were significantly higher than their parental lines in rate of fertilization (t = 7.99, df = 7, p-value < 0.001, F1 hybrids: 72.93 %, parental mean: 11.20 %) and in seeds per pod (t = 7.51, df = 7, p-value < 0.001, F1 hybrids: 3.04, parental mean: 1.25), indicating a relative heterosis of 5.51 for rate of fertilization and 1.43 for seeds per pod. The correlation of both traits is high and significant when all genotypes were included (r = 0.71, p-value <0.001), but medium and not significant (r = 0.39, p-value = 0.17) within the group of inbred lines (Fig. 8).

**Fig. 8.**
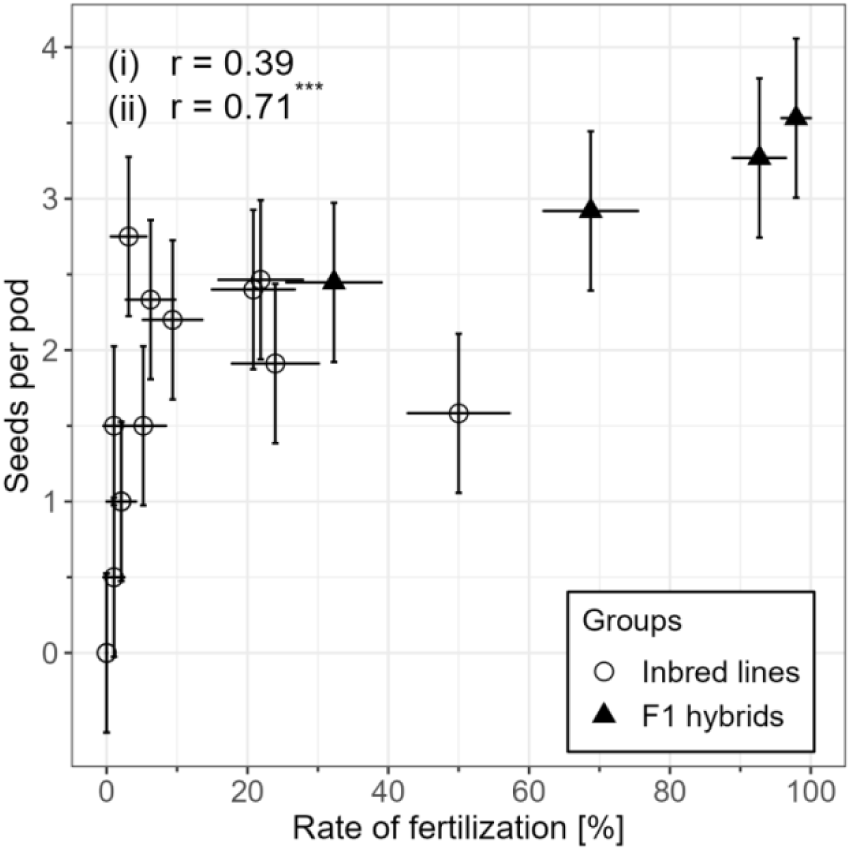
Correlation of rate of fertilization with seeds per pod of four F1 hybrids and 14 inbred lines. Error bars show standard errors of the least square means. Correlation coefficients are calculated based on Pearson’s product moment correlation coefficient for (i) only inbred lines, (ii) all genotypes (*** = significant at p < 0.001)

It can be resumed that there is a trend of autofertile genotypes also having more seeds within the larger number of pods than autosterile genotypes, however, a significant correlation could only be found when F1 hybrids and inbred lines were both included, indicating a heterotic group effect.

### 3.6 Correlation of pollen-related traits

To check whether field data can be reproduced from easier-to-assess pot experiments, we correlated pollen production in the field with pollen production LOW of the pot data. The correlation was low and non-significant for the six inbred lines (r=0.31, p-value=0.56, Spearman’s rank correlation test) and for the six inbred lines and two F1 hybrids combined (r=0.21, p-value=0.62, Spearman’s rank correlation test), indicating an environmental effect on the rank of genotypes regarding their pollen production (Supplementary file 2, Supplementary Fig. 7). Pollen viability was assessed at genotypes out of which only three were included in both the field and the pot experiment, thus, no correlation was feasible.

We then compared our data on autofertility assessed in 2016 to previously published data on autofertility assessed in 2013-2015 (Puspitasari 2017). Rate of fertilization estimated in 2016 was highly and significantly correlated to rate of fertilization estimated in 2013-2015 (r = 0.88, Pearson’s product moment correlation test, t = 5.93, df = 10, p-value < 0.001, see Supplementary file 2, Supplementary Fig. 8). As effect, we used rate of fertilization assessed in 2016 in subsequent correlation tests.

To test the relationships between autofertility and pollen-related traits estimated in the pot experiment, we employed Pearson’s product moment correlation tests (Fig. 9). Correlation coefficients for the group of inbred lines were medium (pollen production) to high (pollen viability), but no significant correlation was found between any of the traits. To resume, we found no evidence for a relationship between pollen production and autofertility. Likewise, no relationship between autofertility and pollen viability was found given the limitations of only four tested inbred lines.

**Fig. 9.**
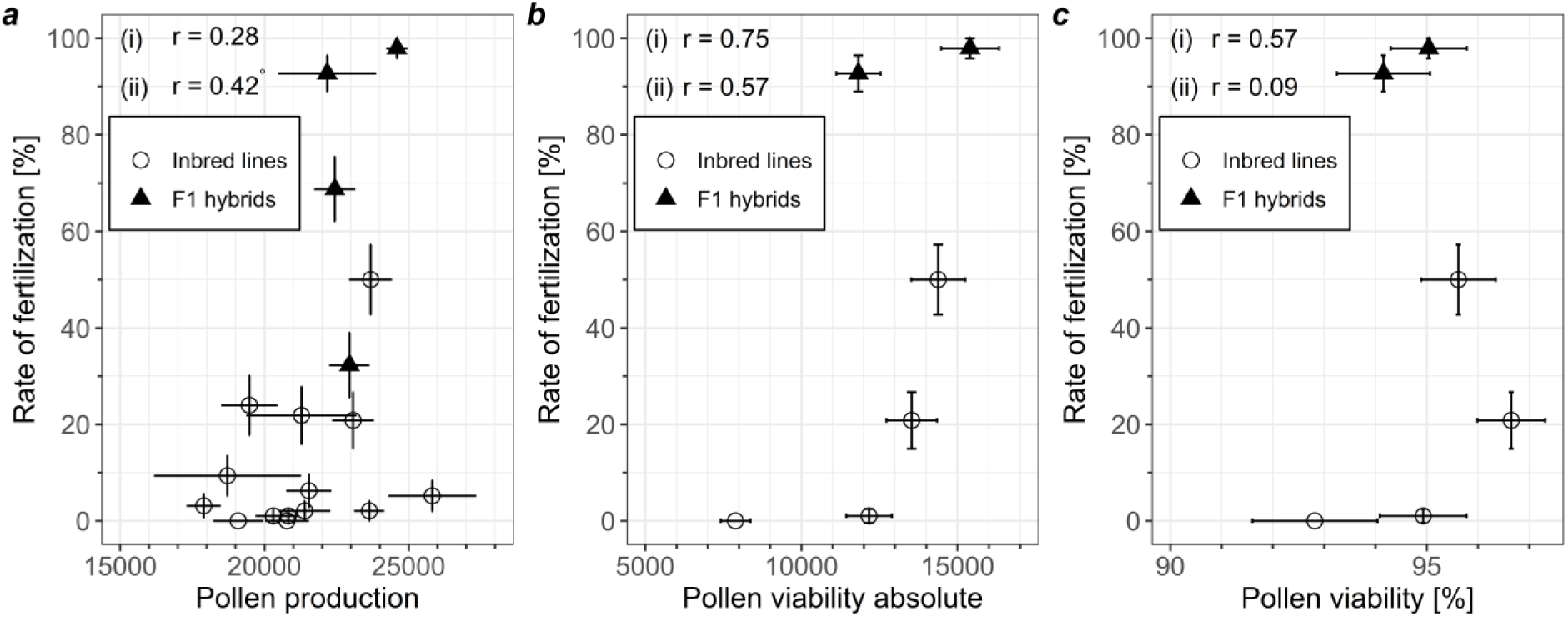
Correlation of rate of fertilization with **a** pollen production LOW, **b** pollen viability absolute, **c** pollen viability [%]. Shown are Pearson’s product moment correlation coefficients for (i) only inbred lines, (ii) all genotypes (° = significant at p = 0.1). Error bars show standard error of the means. All traits were assessed in the caged outdoor pot-experiment 2016

Finally, the relationships between paternal outcrossing success and other pollen-related traits estimated in the field experiment were tested (Table 6). In the six inbred lines, pollen production and absolute number of viable pollen grains were highly correlated to paternal outcrossing success (r=0.77 and r=0.80, respectively, significant at p=0.1), whereas relative pollen viability and rate of fertilization were low to medium and non-significantly correlated to paternal outcrossing success (Supplementary file 2, Supplementary Fig. 9). Interestingly, the degree of cross-fertilization was negatively however non-significantly correlated to pollen production and pollen viability. In sum, pollen production and absolute pollen viability might be positively related to paternal outcrossing success whereas paternal outcrossing success does not seem to be related with neither relative pollen viability nor autofertility.

**Table 6.** Correlations between pollen-related traits estimated with inbred lines (bold number, N=6) and with inbred lines and F1 hybrids (N=8). POS =Paternal outcrossing success, DC = Degree of cross-fertilization. All traits were except rate of fertilization were assessed in the field experiment. Data on POS and DC was taken from Brünjes and Link (2021). °, *, *** denote significances at p= 0.1, p=0.05 and p=0.001, respectively

|  | POS | DC | Pollen production | Pollen viability absolute | Pollen Viability relative | Rate of fertilization |
| --- | --- | --- | --- | --- | --- | --- |
|  | Field data |  |  |  |  | Pot data |
| POS | - | **-0.80\***<br>-0.81* | **0.77°**<br>0.57 | **0.80°**<br>0.63° | **0.32**<br>0.68° | **0.19**<br>0.69° |
| DC | - | - | **-0.33**<br>-0.66° | **-0.39**<br>-0.69° | **-0.30**<br>-0.73* | **0.43**<br>-0.78* |
| Pollen production | - | - | - | **0.99\*\*\***<br>0.99*** | **0.60**<br>0.71° | **0.64**<br>0.76* |
| Pollen viability absolute | - | - | - | - | **0.65**<br>0.76* | **0.58**<br>0.76* |
| Pollen viability relative | - | - | - | - | - | **0.07**<br>0.65° |

## 4. Discussion

### 4.1 Pollen production

#### 4.1.1 Effects of genotype on pollen production

In this study, we report pollen production, i.e. number of pollen grains per flower, in a set of 18 winter faba bean genotypes, both F1 hybrids and inbred lines, assessed with impedance flow cytometry under consideration of the position of the sampled inflorescence. Pollen production in faba bean has not yet been studied extensively, thus, we present the trait in several aspects including heterosis and node position of sampled inflorescence.

Pollen production varied extensively in our set of 18 genotypes, ranging from 17000 to 26000 pollen grains per flower. Previous studies on pollen production in faba bean reported varying numbers of pollen grains per flower, ranging from 1500 (Kambal et al. 1976) to over 50000 (Suso et al. 2008). This large variation might be grounded in the method of pollen counting, the plant material (e.g. inbreeding level, breeding material), and the position of the sampled inflorescence. Most previous studies employed a counting method adapted from Kambal et al. (1976), using a haemocytometer slide (e.g. Carré et al. 1994; Suso et al. 2008; Chen 2009; Bailes et al. 2017), whereas Aguilar-Benitez et al. (2022) used a particle counter. With our method employing an impedance flow cytometer, we obtained pollen counts similar to those found by Carré et al. (1994), Chen (2009) and Aguilar-Benitez et al. (2022), although our numbers were much higher than those reported by Kambal et al. (1976) and lower than those by Suso et al. (2008).

Indeed, we found significant differences between genotypes with regard to pollen production. F1-hybrids showed higher pollen production than inbred lines, reflecting heterosis for this trait. We identified absolute and relative heterosis for pollen production of 1600 pollen grains and 8.6 % at the lower inflorescences and 1900 pollen grains and 10.7 % at the upper inflorescences, respectively, which was significant although the size of heterosis was not very pronounced. When testing each specific cross, only two out of four crosses showed significant heterosis. Similar to the findings of Kambal et al. (1976), in our data, the size and the significance of heterosis for pollen production depended largely on the cross and those inbred lines with the highest pollen production were similar in pollen production to the F1 hybrids. Likewise, for sunflower, Vear et al. (1990) found heterosis for pollen production in ten out of 17 hybrids. We could not assess whether parental lines with high pollen production result in F1 hybrids with high pollen production, as the analysis of the correlation was not feasible with only four F1 hybrids.

Interestingly, pollen production varied less with only four hybrids than with the inbred lines. Whereas some inbred lines had a low pollen production of clearly less than 20000 pollen grains, others such as S_085 and S_199 (over 23000 pollen grains) partly even exceeded the F1 hybrids (Fig. 4). One simplified model describing the occurrence of heterosis is the dominance theory (Crow 1999), assuming that the favorable alleles act dominantly and thus, a heterozygous state in F1 hybrids fully compensates the unfavorable recessive allele. Our findings appear to be in line with such simplified model: S_019 and S_035 carry different, unfavorable alleles in homozygous state, which are compensated for in their heterozygous hybrid, whereas other inbred lines carrying a high share of favorable alleles do not differ much from their hybrid.

#### 4.1.2 Effects of inflorescence position on pollen production

Inflorescence position had an effect on pollen production. We found a continuous decrease in pollen grains per flower from inflorescence 1 (about 20000) to inflorescence 10 (7000), estimated at six genotypes. Likewise, in a further experiment on 18 genotypes, pollen production was higher on the lower part of the tiller (inflorescence 4 or 5) than on the upper part of the tiller (inflorescence 7 or 8) (Fig. 2 and 4). Here, pollen production was higher than in our other experiment where pollen production was estimated together with viability (Table 4) and plants were sampled at higher floral positions (mean: inflorescence 9.45). A similar trend was found by Carré et al. (1994), who counted less pollen grains at inflorescence 10 than at inflorescence 5 in one inbred line, which further supports our findings of lower pollen production at higher floral nodes. A decrease in pollen production per flower over the course of the flowering season has been reported for other plants (Young and Stanton 1990; Lau and Stephenson 1993) due to an increase in prior flowers and seeds. Thus, it is relevant to mention the position of the sampled inflorescence when reporting figures of pollen production in faba bean. In our study, position of the sampled inflorescence and the interaction of inflorescence position with genotype were significant effects for pollen production across inflorescences. However, there was no significance of the differences between genotypes, because we had only two replicates and large genotype-by-replicate interactions (Table 5). To get a significant main effect of genotypes, we would (given all else unchanged) have to increase the number of replicates to four (p=0.009 instead of p=0.321). Some genotypes (e.g. S_019) produced flowers across all ten nodes, whereas others (e.g. F1(S_199xFam157)) only developed six inflorescences and then stopped flowering. If genotypes were a significant main effect on pollen production across inflorescences, it would be favorable to select those genotypes as parental components of a synthetic that produce amounts of pollen that are similar to each other and possess a similar flowering period. Such, the genetic contribution (i.e. genetic dose) of genotypes as fathers in Syn-0 for Syn-1 would ideally not differ between genotypes and consequently, inbreeding in Syn-1 and further generations would be minimized.

#### 4.1.3 Effects of environment on pollen production

Pollen production was higher in potted plants growing in isolation cages than in plants growing in the field, assed at the same genotypes, inflorescence position and year (Supplementary file 2, Supplementary Fig. 7). Concluding from the low and non-significant rank correlation between pot data and field data, the environment seems to have a pronounced effect on pollen production. Part of this environmental effect might be that in the field, plants were non-standardized in terms of number of inflorescences, flowers per inflorescence and number of tillers. Such, plants in the field had to distribute the available resources among a higher number of flowers. Aufhammer and Götz-Lee (1991) reported that removing the three lowest inflorescences in faba bean increased the seed yield at the remaining inflorescences. A similar effect of redistributing reproductive efforts might take place in pollen production, resulting in a lower amount of pollen per single flower in the presence of many flowers. In a study on 21 bee-pollinated legume species, Vonhof and Harder (1995) found that flower production (per inflorescence or per plant) significantly influenced the number of pollen grains produced per flower in only eight species, among them *Vicia americana*. The implications on *Vicia faba* remain unclear, as the effect of standardization treatment on pollen production has not been studied, here.

### 4.2 Pollen viability

#### 4.2.1 Effects of genotype on pollen viability

The overall percentage of viable pollen grains in total pollen grains per flower was very high, both in the field (87.5 % to 94.9 %) and in the pot experiment (92.8 % to 96.6 %). We found significant differences in absolute and relative viability between genotypes. In the field experiment, the two F1 hybrids as a group were significantly higher in absolute pollen production than the group of inbred lines, but several inbred lines were higher in absolute viability than one of the hybrids. Accordingly, absolute viability mostly depended on pollen production, as relative viability was throughout very high. In relative viability, however, the F1 hybrids were consistently higher than any inbred line, suggesting that heterosis might indeed play a role. However, we could not study heterosis of viability in the field trial, as no parental lines were included. In contrast, in the pot experiment, there was no difference in absolute or relative viability between F1 hybrids and their parental inbred lines.

In breeding, the significant differences in absolute and relative viability between genotypes could be considered in the composition of a synthetic. Similar to pollen production, those genotypes should be selected that differ little in viability to facilitate equal doses of every genotype in the Syn-1.

#### 4.2.2 Effects of environment and assessment method on pollen viability

Only in the field experiment, F1 hybrids were significantly higher in relative pollen viability than inbred lines. It thus seems that the uncontrolled conditions in the field led to a certain decrease in relative pollen viability, an effect which F1 hybrids could compensate better than inbred lines. Drought or heat stress during pollen development can reduce pollen viability and pollen germination (Weaver and Timm 1988; Shivanna 1991; Djanaguiraman et al. 2013; Jiang et al. 2015; Bishop et al. 2016; Bheemanahalli et al. 2019). Besides environmental stress, pollen viability and germination can be affected by dehydration during storage (Shivanna and Heslop-Harrison 1981; Brunet et al. 2019), by the method of pollen separation from the flower (Kron et al. 2021) and by the method of viability assessment (Impe et al. 2020). As we analyzed the pollen within few minutes after bud sampling, we reduced any effect of pollen dehydration to a minimum. What is the optimal way to separate pollen from *Vicia faba* flowers to estimate viability? We extracted the intact anthers at one day before dehiscence, which is an easy method to obtain the complete pollen load. However, viability was hence not assessed at a date that reflects the actual conditions for reproduction, as one day before dehiscence, bees have not yet access to the flower and viability is probably not optimal yet. Another possibility is to sample the flower later, at one day before anthesis, just before bees would have access which would be better to reflect real conditions. However, the pollen will already be released from the anthers hence the pollen would have to be gathered from the keel. Such, only the ratio of viable pollen could be assessed, not the total amount of pollen (Bishop et al. 2016). We used impedance flow cytometry (IFC) to assess pollen viability, which proved itself as an efficient and fast method to estimate pollen fertility and to analyze a high number of flower buds directly after sampling. Meanwhile, we want to emphasize that viability is not to be taken synonymously with pollen germination or pollen tube growth. Rather, viability as assessed with IFC is an indicator or a pre-requisite for pollen germination, but not a guarantee (Heidmann et al. 2016). This notion is confirmed by Impe et al. (2020) and Langedijk et al. (2023), who directly compared methods to analyze wheat and Solanaceae pollen, respectively, and indeed found high and significant correlations between the results of in vitro pollen germination and IFC viability, but also higher IFC viability rates than *in vitro* pollen germination rates tested on adapted media. Thus, the high viability we found in faba bean pollen is rather to be taken as predictor for the pollen germination rates of different genotypes, than a direct estimation of pollen germination rates.

The method of viability assessment can affect a study’s finding with regard to how it deals with small pollen grains. A normal-sized faba bean pollen grain has about 50 µm length and 30 µm width (Poulsen and Martin 1977; Supplementary file 1, Fig. 1). Dead pollen grains are smaller than viable ones (Heidmann et al. 2016; Aguilar-Benitez et al. 2022), and if pollen grains of any size are used to calculate viability, the results will be different than if only normal-sized pollen is used. For instance, in their study on faba bean reproductive traits, Aguilar-Benitez et al. (2022) employed one inbred line with a high pollen production, but 70 % of the pollen grains were small in size (i.e., equatorial width lower than 27 µm), resulting in a low viability rate of 26 %. The second inbred line they tested showed viability rates of about 78 %. In contrast, in our study conducted with IFC, we identified viability rates from 88 % to 97 %. This difference may partly result from the different plant material, but may partly be explained by the difference in methods. In our study, small pollen grains may either have not been detected due to too highly set trigger values (Amphasys AG 2023, personal communication) or deformed non-viable pollen may have been discarded as debris, thus resulting in a lower number of counted non-viable pollen grains and overall higher viability rates. In IFC, the amplitude provides information about cell size (Heidmann et al. 2016), however, the IFC data allows only to compare the relative size pollen grains within a sample, but not a direct measurement of size.

Concluding, results on pollen production and pollen viability partly depend on the assessment method (e.g., IFC, particle counter, haemocytometer slides) and limit a direct comparison of absolute numbers between different studies. However, within a study, any technique allows to identify the relative differences between genotypes.

### 4.3 Autofertility

Autofertility varied in inbred lines from 0.0 % to 50.0 % (mean: 10.5 %) and in F1 hybrids from 32.3 % to 97.9 % (mean: 72.9 %) thus almost fully exploiting the quantitative variation of this trait from (0 % to 100 %, Link et al. 1994a). Just as other studies (Drayner 1959; Stoddard 1986; Link et al. 1994a), we identified heterosis for autofertility. Hybrids were significantly higher in rate of fertilization than their parental lines (t-test: t = 7.99, df = 7, p-value < 0.001) and expressed a high value for relative heterosis of 5.51, thus a more than fivefold higher value in F1 compared to their inbred parents. Autofertility has been described as highly heritable trait (Link 1990). Our data confirms a high heritability in the broad sense, i.e., a high reproducibility of the results in different environments by comparing our autofertility data of 2016 to the results of 2013-2015 (Puspitasari 2017).

How is autofertility related to pollen production – Does high pollen production result in high autofertility? Our study indicates that high pollen production rather not leads to higher autofertility. We found a low positive correlation between pollen production and autofertility, which was only significant at p=0.1 when the heterotic effect of autofertility was included in the data (Fig. 9a). Generally, we found that the four tested F1 hybrids were high in autofertility and average in pollen production. Four out of 14 inbred lines (S_025, S_085, S_199, S_145) were similar or higher in pollen production LOW than the four F1 hybrids (Fig. 9a). Previous studies likewise found no consistent trend between autofertility and pollen production. Kambal et al. (1976) noted that genotypes high in pollen production were not necessarily high in autofertility, rather, they identified an association between high pollen production and low autofertility. Similarly, Aguilar-Benitez et al. (2022) counted double as many pollen grains in an autosterile line than in an autofertile line, but, in contrast, when only considering normal-sized pollen, the autofertile line showed a higher pollen production than the autosterile line. Alike the latter finding, Chen (2009) identified higher pollen production in autofertile F1 hybrids and in their autofertile F2 and F3 segregants than in the autosterile lines. We conclude that, although both traits, autofertility and pollen production, are rather high in F1 hybrids, this is probably not a matter of cause and effect because both traits are decoupled in inbred lines. A clear relationship between both traits does not seem to be the case.

Autofertility and absolute or relative pollen viability were weakly correlated in the field (N=6) and medium-to-high but non-significantly in the pot experiment (N=4). Given the low number of tested genotypes, correlation tests of autofertility with pollen viability might become significant for a larger sample size. However, our data does not strongly imply a relationship between both traits.

The connection between autofertility and degree of cross-fertilization or paternal outcrossing success is less obvious. Highly autofertile genotypes might self-fertilize already before anthesis so that cross-pollen transported by bees arrives too late as ovules are already fertilized (Poulsen 1980). A similar biological connection might hold between autofertility and paternal outcrossing success: if ovules are already self-fertilized, fewer options to realize paternal outcrossing success will exist. This notion was supported by Link (1990) who found a strong and highly significant negative correlation between autofertility and degree of cross-fertilization in a group of F1 genotypes, however, only a low and non-significant negative correlation in a small set of nine parental inbred lines. Our set of six inbred lines and two F1 hybrids likewise was too small to show a significant relationship between autofertility and degree of cross-fertilization or paternal outcrossing success. Further, although the genotypes of the six inbred lines tested in the field experiment had been chosen to cover a wide range of autofertility levels (Table 1), we found all lines except one to be low in autofertility (Supplementary file 2, Supplementary Fig. 9d). To improve the explanatory power of correlation studies involving autofertility, a higher number of genotypes with varying levels of autofertility should be tested.

### 4.4. Paternal outcrossing success

Since we found no clear relationship between autofertility and pollen production (N=18, see Fig. 9a), we conclude that autofertility does not affect paternal outcrossing success via pollen production. Paternal outcrossing success was estimated for eight genotypes, whereas autofertility was assessed at 18 genotypes. Why couldn’t we analyze the correlation between autofertility and paternal outcrossing success for all N=18 genotypes? The reason was not lack of funding but rather the limitations on how to estimate paternal outcrossing success. To identify the father genotype with a limited number of SNP markers (see Brünjes and Link 2021), the number of genotypes per polycross cannot be increased arbitrarily, and as the values for paternal outcrossing success depend on each specific set of genotypes, we cannot merge the values from several polycrosses together for analyses. Thus, the number of evaluated genotypes per analysis is limited, here limited to N=8, making it more difficult to obtain significant correlations of paternal outcrossing success with other traits.

Pollen production and absolute pollen viability were highly correlated with paternal outcrossing success, significant at p=0.1. Our data indicates that it is more likely that pollen production and absolute viability are connected to paternal mating success than that they are not. The correlation of relative viability with paternal outcrossing success was low and non-significant which, however, not necessarily objects any relationship between these traits. In the field trial, we had little variation in relative pollen viability of the inbred lines, ranging from 91.3 % to 93 %, except for one line that was slightly lower. Given such overall high viability rates and overall high pollen production, we conclude that the paternal outcrossing success of the tested genotypes will most likely not be impeded by limited pollen quantity or quality.

While it seems plausible at first glance that high pollen production results in high paternal outcrossing success, the biological connections are complex and multilayered. Whereas pollen production sets the upper limit of a plant’s potential paternal success, the pathway from the pollen-producing father plant to a healthy developed seed growing on the mother plant comes with multiple divergences where pollen may get lost (Minaar et al. 2019). For example, pollen production per flower differs between species with different type of pollen dispersal. In self-fertilized peanut (*Arachis hypogaea*), most pollen will be retained within the flower, and pollen production is relatively low (about 8000 pollen grains per flower, Prasad et al. 1999). In contrast, wind-pollinated species such as maize have been shown to produce pollen on a much higher level (2 Mio. to 4 Mio. pollen grains per plant and day, Uribelarrea et al. 2002). In animal-pollinated crops, pollen production has been shown to lie in between the previous examples. In faba bean, we found 17000 to 26000 pollen grains, which is similar to the pollen production of other insect-pollinated crops such as *Cucurbita pepo* (30000 to 43000 pollen grains per flower, Lau and Stephenson 1993, 1994, 1995; Vidal et al. 2006) and *Helianthus annuus* (33000 to 36000 grains/floret, Vear et al. 1990). An animal-pollinated plant investing in male fitness by increased pollen production needs an increased vector visitation rate or increased placement of pollen loads on vectors as premises to get its pollen transported to other plants’ stigmas – otherwise, its pollen may just be left in the anthers (Minaar et al. 2019). Offering large amounts of pollen as a reward to pollen vectors to support pollen export can also have detrimental effects on reproductive success as bees collect the protein-rich pollen and consume it either directly or to feed their larvae, thereby diminishing the siring opportunities of the plant (Hargreaves et al. 2009). Further, the composition of pollinator communities may affect the link between pollen production and paternal outcrossing success. In wild radish, greater visitation rates by honey bees significantly reduced paternal success whereas plants high in pollen production received more visits from small wild bees relative to honey bees, thereby diminishing the adverse effects of honey bee visits on paternal success (Stanton et al. 1991). In general, where pollinators are diverse and floral resources are limited, a plant that attracts the most efficient pollinator likely exhibits a higher probability of getting its pollen transported to a receptive flower, resulting in higher paternal success. Hence, the realized paternal success is evidently not only depending on pollen production, but also on the interaction between plants and pollen vectors. Even pollen that has successfully been delivered to a stigma does not ensure male reproductive success of the father plant, as there may be more pollen grains deposited on the stigma than there are ovules, and usually pollen grains from more than one donor are present (Cruzan 1990). Competitive abilities of the different pollen donors in postpollination characters such as pollen germination, pollen tube growth or fertilization ability work even beyond pollen transfer (Marshall and Diggle 2001). To study which of these developmental mechanisms is associated with paternal outcrossing success and how pollen donors might interact would be a further research question.

Until the widespread use of genetic markers, direct information on paternity was difficult to obtain and until today, only few studies have empirically tested the relationship between pollen production and paternal success. The conclusions of the limited number of studies are not straightforward. In wind-pollinated white spruce trees, an increased cone production led to an increase in paternal success (Schoen and Stewart 1986). In insect-pollinated wild radish (*Raphanus sativus* L.), pollen production of the donor plant had a significant effect on the realized paternal success. Stanton et al. (1991) found that radish plants with greater pollen production sired more seeds, however, this was an indirect effect as plants high in pollen production also attracted more small bees than their competitors, thereby diminishing the detrimental effects of honey bee visits on paternal success. Marshall (2007) concluded that apart from differences in pollen production, also the pollen donor identity significantly affected paternal success in wild radish, thus paternal characteristics other than pollen production likely play a role. Surprisingly, Ashman (1998) found no significant link between differences in pollen production and paternal success in wild strawberry (*Fragaria virginiana*), instead, the amount of pollen removed from a flower significantly correlated with paternal success. Thus, the potential for pollen production to influence paternal success directly may be limited by the plant-pollinator interplay, which is a fundamental link between both characteristics (Minaar et al. 2019).

Apart from the ambiguous connection between pollen production and paternal success, pollen production itself is highly variable within and between species, and a large part of this variation can be explained by environmental factors (Minaar et al. 2019). However, Young et al. (1994) found pollen production in wild radish to be highly heritable and genotype-environment interactions were non-significant. In our study, the ranks of genotypes changed in both environments due to genotype × environment interaction. Hence, different genotypes would be selected in each environment. Based on our data, pollen production in faba bean is indeed an environment-specific trait, rendering it necessary to select in the environment where seed production with the thus-selected genotypes shall take place.

### 4.5 Relevance for plant breeding

The significant differences between genotypes we found imply that it would be possible to select genotypes based on their pollen characteristics. Sufficient pollen production and pollen viability are important for the success of genotypes as male parent and, consequently, those genotypes that are markedly low in these characters could be avoided. Furthermore, to compose synthetics, the aim is equal doses of paternal contribution of each component to the generation Syn-1, thus, components that strongly differ in pollen production, viability or paternal outcrossing success should be avoided, to minimize inbreeding in later generations. Accordingly, parental components should be chosen based on similar instead of highest values, i.e. those genotypes that, as a group, show minimum differences in these traits. Such, selection would be performed in a stabilizing manner.

Assessing paternal outcrossing success is costly and not feasible to implement in the breeding process. In contrast, assessing pollen production and viability of candidate genotypes is easy and fast to accomplish when using a cell counter or impedance flow cytometer. Our results suggest that pollen production is highly related to paternal mating success, although at a weak significance level. Thus, characterizing the genotypes in their pollen characteristics might be a worthwhile step as this information could be used when composing the Syn-0 where the set of components should be similar in pollen characteristics.

Pollen production per flower clearly decreased with the progress of flowering across inflorescences, thus, our results stress the importance of equal begin of flowering in the chosen components. Faba bean genotypes can bear over ten nodes of inflorescences (Link and Stützel 1995), but mostly only the lower inflorescences develop pods. In a scenario where components are not synchronous in their flowering period, earlier-flowering genotypes presumably have a reduced chance to bear seeds sired by late-flowering genotypes. Once the late-flowering genotypes are in full bloom and highly attractive to pollinators, the earlier-flowering genotypes meanwhile blossom at their upper inflorescences that produce less pollen and lower number of flowers, thus exhibiting a reduced attractiveness to pollinators (Suso and Maalouf 2010). Hence, asynchronous flowering among parental components probably reduces inter-genotype cross-fertilization and increases inbreeding in the subsequent generation.

Pollen number and quality is not only of interest in faba bean breeding, but in breeding and seed production in general. First, sufficient quantity and quality of pollen is required as primary prerequisite to ensure male reproductive success. Second, differences in pollen quantity and quality among genotypes may result in differences in paternal outcrossing success which influence both the level of inbreeding and allele frequencies in subsequent generations (Brünjes and Link 2021). Thus, considering pollen properties might not only be interesting in faba bean, but also for breeding of synthetics of partially allogamous crops with a marked degree of cross-fertilization and even more so in completely allogamous crops.

## 5. Conclusions

In this set of experiments, we checked the relationships between pollen production, pollen viability, autofertility and paternal outcrossing success in winter faba bean. We found significant differences in both pollen production and pollen viability, but the size of the differences was not very pronounced and the amount and quality of pollen was overall high. We conclude that pollen quantity and quality in faba bean is probably sufficient to ensure male reproductive success. Further, we aimed to find out whether pollen characteristics and autofertility can be used as auxiliary traits to estimate paternal outcrossing success. Whereas we found no evidence for a direct relationship of autofertility and pollen production, pollen quality or paternal outcrossing success, pollen production and the absolute number of viable pollen grains were highly correlated with paternal outcrossing success. When selecting inbred lines as polycross components, assessing pollen characteristics might be a useful tool to reduce differences between genotypes in paternal mating successes, thereby minimizing inbreeding.

## Acknowledgements

We thank Anna Pfeiffenschneider for taking care of plants in the pot experiment and her highly valuable contribution in sample analysis with the impedance flow cytometer. Winda Puspitasari standardized plants of the pot experiment and kindly provided us with her autofertility data from previous years. We further thank Regina Martsch and Andreas Henn for their indispensable support in implementing experiments and caring for the plants. We like to thank Philipp Körner and Marco Di Berardino from Amphasys AG for their technical support and helpful advice. The authors acknowledge the German Federal Environmental Foundation (Deutsche Bundesstiftung Umwelt, DBU, AZ 20014/302) for funding.

