## Supplementary File 1 for "Pollen production, pollen viability and autofertility in faba bean (*Vicia faba* L.) and their relationship with realized paternal success"

**Testing faba bean with impedance flow cytometry**

To prepare the samples for analysis, the ten closed anthers of a flower bud were transferred into an Eppendorf tube and softly squeezed with a pistil to release the pollen grains from the anthers (Fig. 1a). Then, 1 ml of buffer solution (AF 5, Amphasys, Lucerne, Switzerland) was added by rinsing adhering pollen grains off the pistil into the Eppendorf tube. After homogenizing the sample for 20 sec, the buffer with the pollen was filtered through a 100 µm sieving filter into a second Eppendorf tube. Another 1 ml buffer was added to the first Eppendorf tube, homogenized, and filtered through the same filter, thereby rinsing the filter and adding any remaining pollen to the buffer mix in the second Eppendorf tube.

After homogenizing the total sample, the solution was loaded into the impedance flow cytometer type Ampha Z30 (Amphasys, Lucerne, Switzerland) while manually shaking the tube to prevent settling of pollen grains at the bottom of the tube. A 120 µm impedance chip was used for measurements. Pollen production and pollen viability were measured at 0.5 MHz employing the default settings (Trigger level 0.1 V, Modulation 3, Amplifier 6, Demodulation 2), while the pump was set to 300 rpm.

Plots produced by the AmphaSoft software (version 1.2.4, Amphasys, Lucerne, Switzerland) displayed different groups of anther debris, non-viable and viable pollen, along with the respective densities, counts and percentages (Fig. 1b). Each dot in Fig. 1b represents one accepted count by the cytometer.

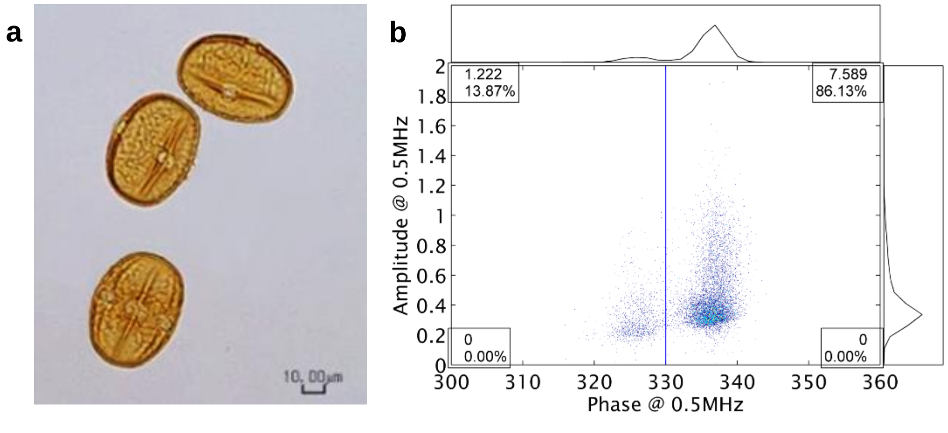


**Fig. 1** Faba bean pollen grains (photo by B. Marzinzig) (**a**) and cytometer dotplot at 0.5 MHz for cell count and viability estimation (**b**)

The software includes a gating function that allows the automatic calculation of the ratio of viable and non-viable pollen. The separation of these two fractions is indicated by a vertical line (line gating), with the viable pollen on the right and the non-viable pollen on the left side of the line.

The optimal line gating for faba bean had not been tested before. Thus, we conducted tests to (i) check the default differentiation between viable and non-viable pollen, and (ii) to determine the optimal line gating for faba bean. To the first aim, we compared fresh, mature pollen with heat-treated aliquots (see Heitmann et al. 2016). The samples were prepared as described above, but only entered with about half their volume into the cytometer and analyzed. The aliquots containing the remaining half of the pollen suspension were exposed to boiling water for 3 minutes, cooled down to ambient temperature and then analyzed. We found that the fresh-pollen samples resulted in high phase angles (Fig. 2a), whereas the heat-treated pollen samples resulted in clearly lower phase angles (Fig. 2b). As the fresh-pollen samples resulted in high phase angles for almost 100 % of the counted pollen grains (Fig. 2a), the optimal line gating for faba bean could not be assigned. Thus, we did further tests with slightly wilted flower buds for which we assumed a higher variation in pollen viability. Flower buds from different genotypes were picked, anthers collected and left to dry for one hour. The cytometer analysis resulted in a clear differentiation between two groups of pollen grains at a phase angle of 330 (Fig. 1b). We concluded that the pollen grains with the higher phase angle in Fig. 2a were the viable ones because they were absent after heat treatment (Fig. 2b). Accordingly, the line gating was set to 330 phase angle and used as template in all samples.

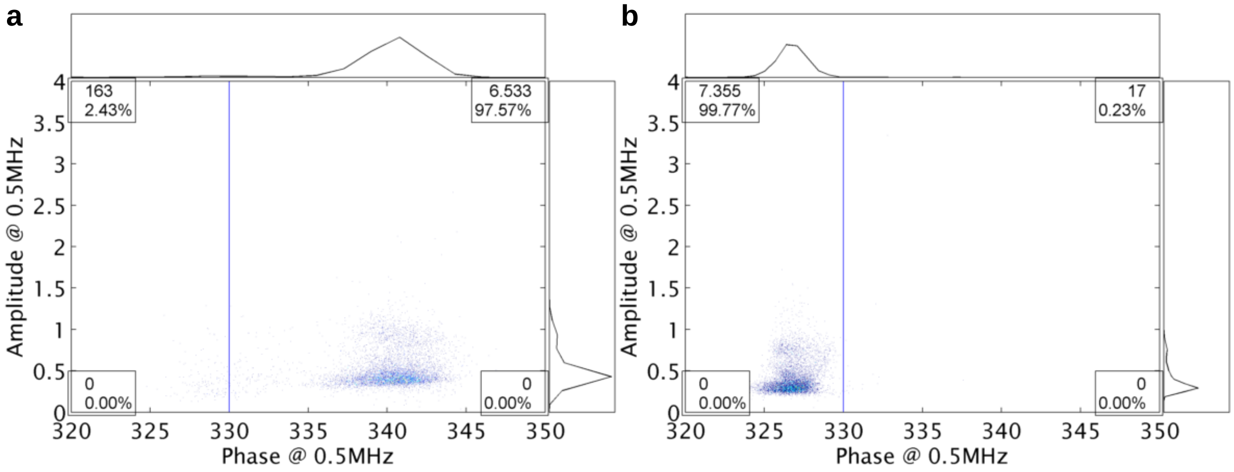


**Fig. 2** Cytometer dotplots at 0.5 MHz of **a** fresh pollen and **b** heat-treated pollen

Estimates for pollen production (i.e., number of pollen grains per flower) were the number of accepted counts. Pollen viability was estimated as (i) the absolute number of viable pollen grains per flower, using the counts of viable cells, and (ii) the percentage of viable cells in total number of accepted counts.
