## Supplementary File 2 for "Pollen production, pollen viability and autofertility in faba bean (*Vicia faba* L.) and their relationship with realized paternal success"

**Supplementary tables and figures**

**Supplementary Table 1** Pollen production, pollen viability absolute and viability relative of eight genotypes in the field experiment. Least square means across 12 replicates. UCI = Upper 95 % confidence interval, LCI = Lower 95 % confidence interval

| **Genotype** | **Pollen production** | **UCI LCI** | **Viability absolute** | **UCI LCI** | **Viability relative [%]** | **UCI LCI** |
|---|---|---|---|---|---|---|
| F1(S_019xS_035) | 16600 | 18640<br>14480 | 15700 | 18100<br>13370 | 94.88 | 97.76<br>92.01 |
| F1(S_025xS_217) | 20200 | 22260<br>18090 | 19100 | 21420<br>16690 | 94.88 | 97.76<br>92.01 |
| S_046 | 16800 | 18890<br>14720 | 15400 | 17770<br>13040 | 91.41 | 94.29<br>88.53 |
| S_085 | 18500 | 20620<br>16450 | 17100 | 19500<br>14780 | 91.68 | 94.55<br>88.80 |
| S_145 | 15000 | 17130<br>12970 | 14000 | 16410<br>11690 | 93.00 | 95.88<br>90.12 |
| S_199 | 15300 | 17390<br>13230 | 14100 | 16430<br>11700 | 91.30 | 94.18<br>88.42 |
| S_235 | 14800 | 16840<br>12680 | 12700 | 15050<br>10330 | 92.53 | 95.40<br>89.65 |
| Fam157 | 12000 | 14100<br>9940 | 9800 | 12180<br>7450 | 87.46 | 90.34<br>84.58 |

**Supplementary Table 2** Mean and standard error of the mean of pollen production (number of pollen grains per flower) at inflorescences 4 and 5 (LOW), inflorescences 7 and 8 (UP) and total plant (TOTAL) of the 18 faba bean genotypes in the pot experiment

| | **LOW** | | **UP** | | **TOTAL** | |
|---|---|---|---|---|---|---|
| **Genotype** | **Pollen production** | **± SE** | **Pollen production** | **± SE** | **Pollen production** | **± SE** |
| **F1(S_019xS_035)** | 22400 | ± 715 | 21100 | ± 571 | 21800 | ± 575 |
| S_019 | 18700 | ± 2545 | 16600 | ± 2019 | 17700 | ± 2255 |
| S_035 | 19100 | ± 858 | 17700 | ± 804 | 18400 | ± 148 |
| **F1(S_025xS_217)** | 22200 | ± 1704 | 22500 | ± 766 | 22300 | ± 1028 |
| S_025 | 23100 | ± 727 | 20400 | ± 1421 | 21700 | ± 1073 |
| S_217 | 20800 | ± 765 | 20600 | ± 1424 | 20700 | ± 603 |
| **F1(S_046xS_085)** | 24600 | ± 369 | 23000 | ± 828 | 23800 | ± 498 |
| S_046 | 20800 | ± 345 | 20200 | ± 985 | 20500 | ± 571 |
| S_085 | 23700 | ± 744 | 21100 | ± 399 | 22400 | ± 534 |
| **F1(S_199xFam157)** | 22900 | ± 700 | 20000 | ± 442 | 21500 | ± 310 |
| S_199 | 23600 | ± 521 | 21200 | ± 208 | 22400 | ± 337 |
| Fam157 | 21500 | ± 781 | 20100 | ± 363 | 20800 | ± 558 |
| S_003 | 20300 | ± 613 | 19700 | ± 1141 | 20000 | ± 660 |
| S_120 | 21300 | ± 1915 | 18400 | ± 208 | 19900 | ± 1003 |
| S_145 | 25800 | ± 1529 | 23900 | ± 783 | 24900 | ± 548 |
| S_235 | 17900 | ± 591 | 18100 | ± 872 | 18000 | ± 656 |
| Diana07 | 21400 | ± 895 | 21800 | ± 1168 | 21600 | ± 983 |
| VF172 | 19500 | ± 973 | 20000 | ± 269 | 19800 | ± 446 |
| **Mean** F1 hybrids | 23000 | ± 542 | 21700 | ± 675 | 22300 | ± 542 |
| **Mean** Parental inbred lines | 21400 | ± 685 | 19700 | ± 593 | 20600 | ± 619 |
| **Mean** All 14 inbred lines | 21300 | ± 587 | 20000 | ± 497 | 20600 | ± 519 |

**Supplementary Table 3** ANOVA table of **pollen production LOW**, estimated with the boxcox-transformed values

| Source of variance | d.f. | MS | F-value | Variance components |
|---|---|---|---|---|
| **Genotypes** | 17 | $147 \times 10^{12}$ | 5.43 | $30 \times 10^{12}$ |
| **Replicates** | 3 | $233 \times 10^{12}$ | 8.61 | $11 \times 10^{12}$ |
| **Error** | 50 | $27 \times 10^{12}$ | | $27 \times 10^{12}$ |
| **Total** | 70 | | | |

**Supplementary Table 4** ANOVA table of **pollen production UP**, estimated with the boxcox-transformed values

| Source of variance | d.f. | MS | F-value | Variance components |
|---|---|---|---|---|
| **Genotypes** | 17 | $5554 \times 10^{12}$ | 3.33 | $972 \times 10^{12}$ |
| **Replicates** | 3 | $1167 \times 10^{12}$ | 0.70 | $-28 \times 10^{12}$ |
| **Error** | 42 | $1667 \times 10^{12}$ | | $1667 \times 10^{12}$ |
| **Total** | 62 | | | |

**Supplementary Table 5** ANOVA table of **pollen production TOTAL**, estimated with the boxcox-transformed values

| Source of variance | d.f. | MS | F-value | Variance components |
|---|---|---|---|---|
| **Genotypes** | 17 | $6200 \times 10^{12}$ | 5.25 | $1255 \times 10^{12}$ |
| **Replicates** | 3 | $2915 \times 10^{12}$ | 2.47 | $96 \times 10^{12}$ |
| **Error** | 41 | $1181 \times 10^{12}$ | | $1181 \times 10^{12}$ |
| **Total** | 61 | | | |

**Supplementary Table 6** Analysis of variance of **absolute heterosis** (absolute pollen number superiority of F1 over the mean of parental inbred lines) for **pollen production LOW**, estimated with boxcox-transformed values

| Source of variance | d.f. | SS | MS | F-value | p-value |
|---|---|---|---|---|---|
| Cross | 3 | $2.14 \times 10^{14}$ | $0.7 \times 10^{14}$ | 2.642 | 0.113 |
| Replicate | 3 | $3.22 \times 10^{14}$ | $1.1 \times 10^{14}$ | | |
| Error | 9 | $2.43 \times 10^{14}$ | $0.3 \times 10^{14}$ | | |

**Supplementary Table 7** Analysis of variance of **absolute heterosis** (absolute pollen number superiority of F1 over the mean of parental inbred lines) for **pollen production UP**, estimated with boxcox-transformed values

| Source of variance | d.f. | SS | MS | F-value | p-value |
|---|---|---|---|---|---|
| Cross | 3 | $153 \times 10^{14}$ | $51.1 \times 10^{14}$ | 8.169 | 0.006 |
| Replicate | 3 | $123 \times 10^{14}$ | $40.9 \times 10^{14}$ | | |
| Error | 9 | $56 \times 10^{14}$ | $6.3 \times 10^{14}$ | | |

**Supplementary Table 8** Analysis of variance of **absolute heterosis** (absolute pollen number superiority of F1 over the mean of parental inbred lines) for **pollen production TOTAL**, estimated with boxcox-transformed values

| Source of variance | d.f. | MS | F-value | p-value |
|---|---|---|---|---|
| Cross | 3 | $41.6 * 10^{14}$ | 4.86 | 0.028 |
| Replicate | 3 | $45.1 * 10^{14}$ | | |
| Error | 9 | $8.6 * 10^{14}$ | | |

**Supplementary Table 9** Analysis of variance of **relative heterosis** (percent superiority of F1 over the mean of parental inbred lines) for **pollen production LOW**, estimated with original count values

| Source of variance | d.f. | SS | MS | F-value | p-value |
|---|---|---|---|---|---|
| Cross | 3 | 1024 | 341.3 | 2.888 | 0.095 |
| Replicate | 3 | 1140 | 380.1 | | |
| Error | 9 | 1063 | 118.1 | | |

**Supplementary Table 10** Analysis of variance of **relative heterosis** (percent superiority of F1 over the mean of parental inbred lines) for **pollen production UP**, estimated with original count values

| Source of variance | d.f. | SS | MS | F-value | p-value |
|---|---|---|---|---|---|
| Cross | 3 | 1450 | 483.2 | 9.471 | 0.004 |
| Replicate | 3 | 898 | 299.3 | | |
| Error | 9 | 459 | 51.0 | | |

**Supplementary Table 11** Analysis of variance of **relative heterosis** (percent superiority of F1 over the mean of parental inbred lines) for **pollen production TOTAL**, estimated with original count values

| Source of variance | d.f. | SS | MS | F-value | p-value |
|---|---|---|---|---|---|
| Cross | 3 | 1122 | 373.9 | 4.916 | 0.027 |
| Replicate | 3 | 976 | 325.4 | | |
| Error | 9 | 684 | 76.0 | | |

**Supplementary Table 12 Absolute heterosis** (absolute pollen number superiority of F1 over the mean of parental inbred lines) for pollen production LOW, UP and TOTAL **in two crosses**, tested with Welch's t-test

| | F1(19x35) and parental lines | | | F1(46x85) and parental lines | | |
|---|---|---|---|---|---|---|
| | t-value | d.f. | p-value | t-value | d.f. | p-value |
| LOW | 2.33 | 5.01 | 0.034 | 4.02 | 5.94 | 0.004 |
| UP | 3.75 | 5.75 | 0.005 | 2.42 | 3.96 | 0.037 |
| TOTAL | 3.14 | 5.16 | 0.012 | 4.07 | 3.80 | 0.008 |

**Supplementary Table 13** Autofertility-related traits of the 18 genotypes. UCI = Upper 95 % confidence interval, LCI = Lower 95 % confidence interval

| Genotype | Rate of fertilization [%] | UCI<br>LCI | Seeds per pod | UCI<br>LCI |
|---|---|---|---|---|
| **F1(S_019xS_035)** | **68.75** | **84.81**<br>**46.43** | **2.92** | **4.76**<br>**1.08** |
| S_019 | 9.37 | 31.27<br>2.30 | 2.20 | 4.04<br>0.36 |
| S_035 | 0.00 | 100.00<br>0.00 | 0.00 | 1.84<br>-1.84 |
| **F1(S_025xS_217)** | **92.71** | **98.53**<br>**70.73** | **3.27** | **5.11**<br>**1.43** |
| S_025 | 20.83 | 43.24<br>8.33 | 2.40 | 4.24<br>0.56 |
| S_217 | 0.00 | 100.00<br>0.00 | 0.00 | 1.84<br>-1.84 |
| **F1(S_046xS_085)** | **97.92** | **99.90**<br>**69.58** | **3.53** | **5.37**<br>**1.69** |
| S_046 | 1.04 | 42.52<br>0.02 | 0.50 | 2.34<br>-1.34 |
| S_085 | 50.00 | 70.34<br>29.66 | 1.58 | 3.42<br>-0.25 |
| **F1(S_199xFam157)** | **32.29** | **54.56**<br>**15.93** | **2.45** | **4.28**<br>**0.61** |
| S_199 | 2.08 | 30.42<br>0.10 | 1.00 | 2.84<br>-0.84 |
| Fam157 | 6.25 | 28.41<br>1.11 | 2.33 | 4.17<br>0.50 |
| S_003 | 1.04 | 42.52<br>0.02 | 1.50 | 3.34<br>-0.34 |
| S_120 | 21.87 | 44.31<br>8.97 | 2.46 | 4.30<br>0.63 |
| S_145 | 5.21 | 27.72<br>0.78 | 1.50 | 3.34<br>-0.34 |
| S_235 | 3.12 | 27.84<br>0.27 | 2.75 | 4.59<br>0.91 |
| Diana07 | 2.08 | 30.42<br>0.10 | 1.00 | 2.84<br>-0.84 |
| VF172 | 23.96 | 46.42<br>10.28 | 1.91 | 3.75<br>0.07 |

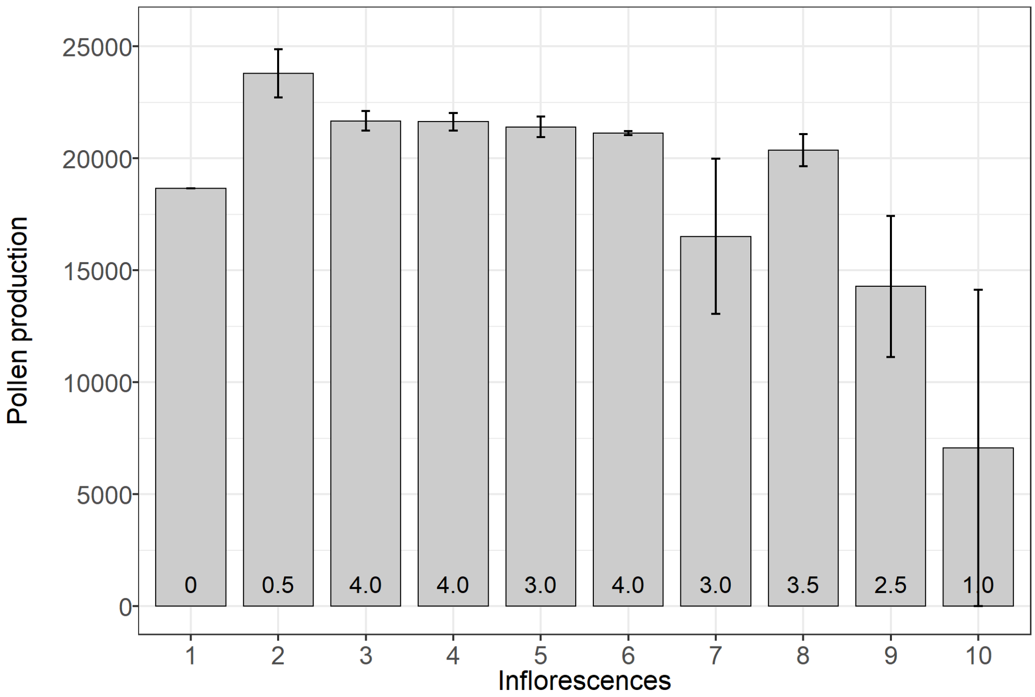

**Supplementary Fig. 1** Pollen production (grains per flower) per inflorescence at the hybrid F1(S_019xS_035).Vertical bars show standard error of the mean across two replicates. Numbers within the bars show the mean number of buds per replicate that entered the analysis

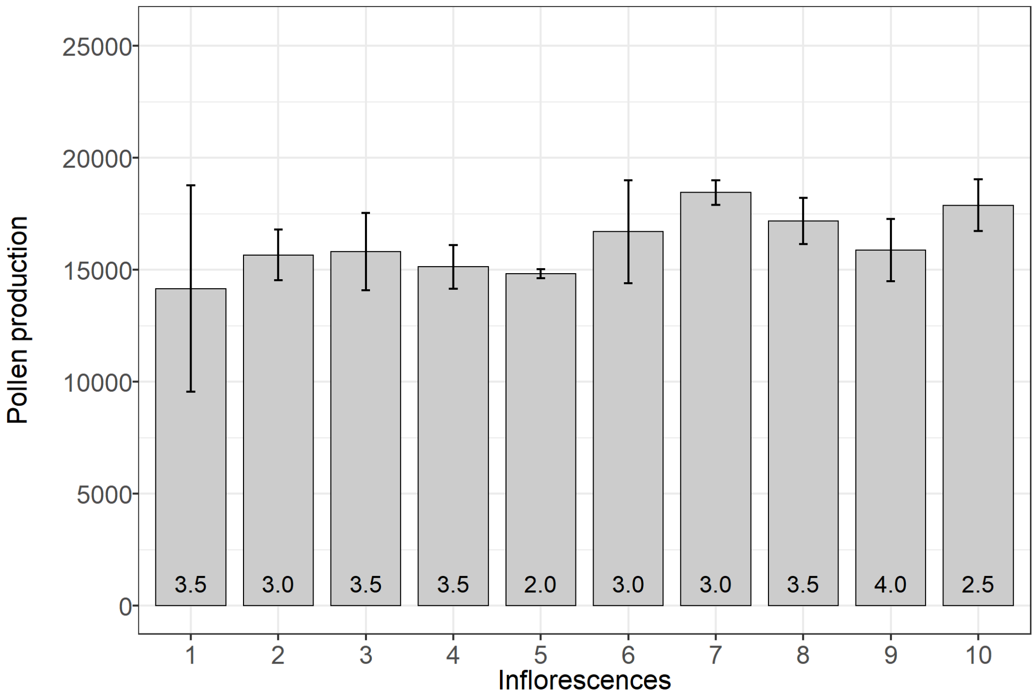

**Supplementary Fig. 2** Pollen production (grains per flower) per inflorescence at the inbred line S_019. Vertical bars show standard error of the mean across two replicates. Numbers within the bars show the mean number of buds per replicate that entered the analysis

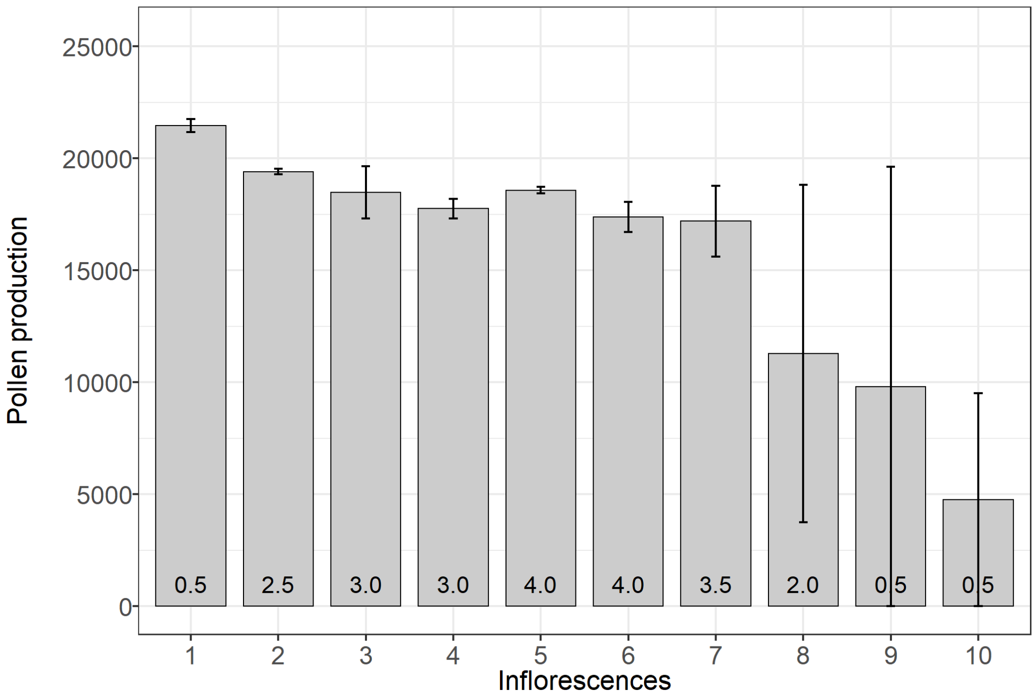

**Supplementary Fig. 3** Pollen production (grains per flower) per inflorescence at the inbred line S_035.Vertical bars show standard error of the mean across two replicates. Numbers within the bars show the mean number of buds per replicate that entered the analysis

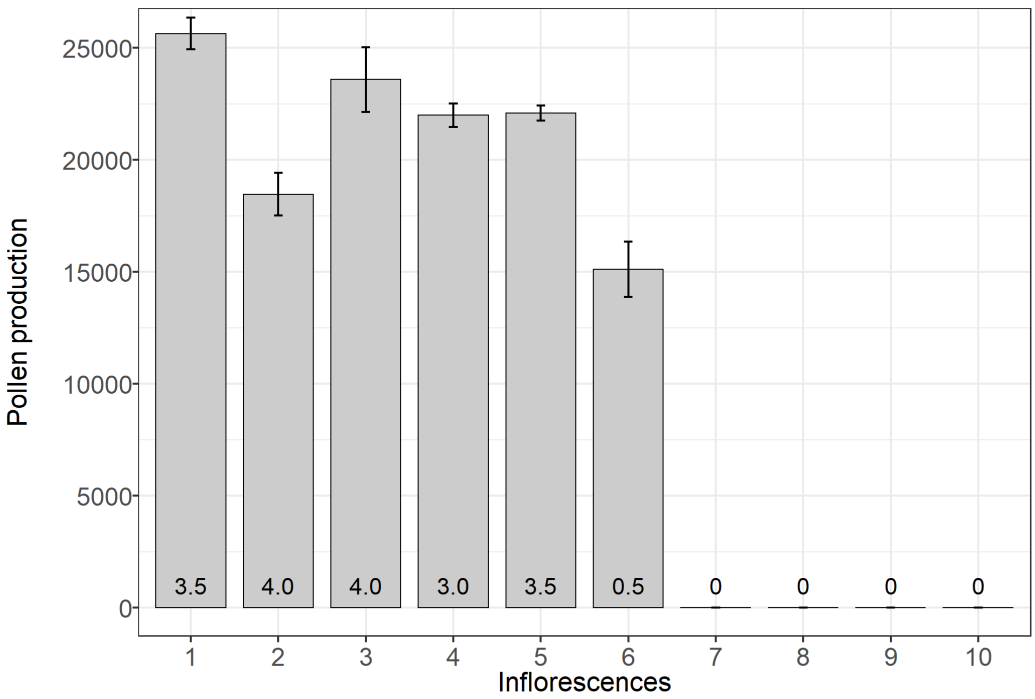

**Supplementary Fig. 4** Pollen production (grains per flower) per inflorescence at the hybrid F1(S_199 x Fam157).Vertical bars show standard error of the mean across two replicates. Numbers within the bars show the mean number of buds per replicate that entered the analysis

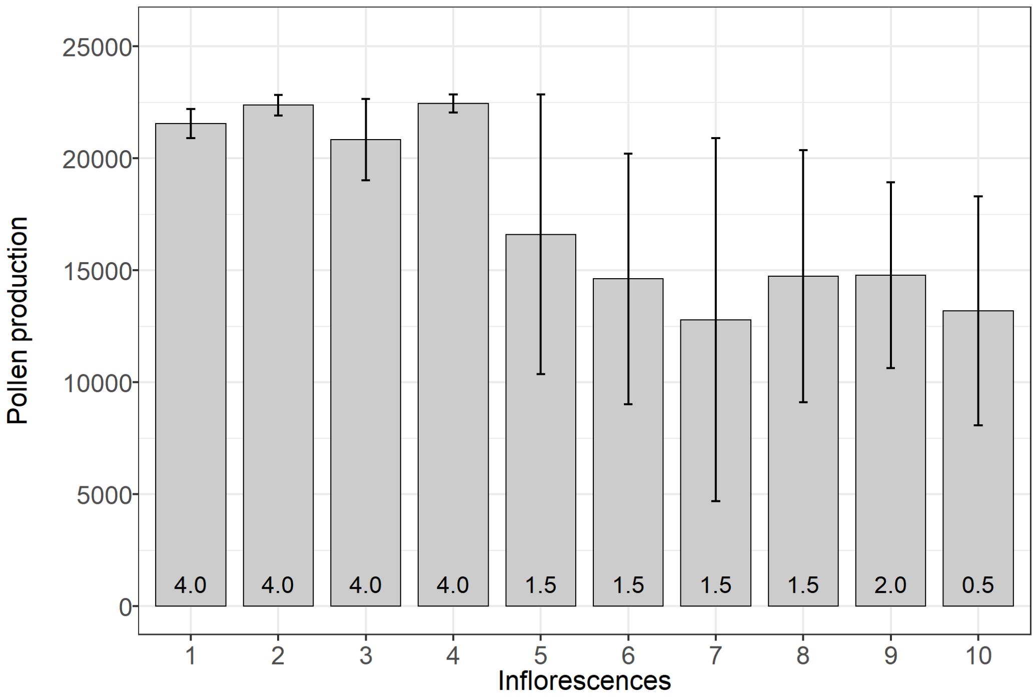

**Supplementary Fig. 5** Pollen production (grains per flower) per inflorescence at the inbred line S_199.Vertical bars show standard error of the mean across two replicates. Numbers within the bars show the mean number of buds per replicate that entered the analysis

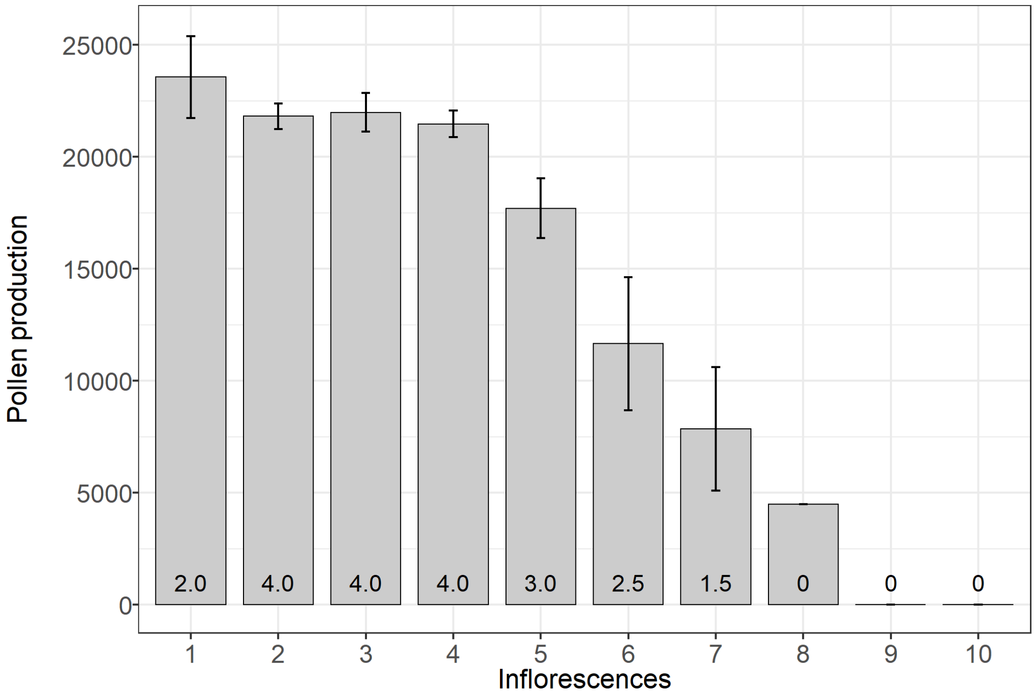

**Supplementary Fig. 6** Pollen production (grains per flower) per inflorescence at the inbred line Fam157.Vertical bars show standard error of the mean across two replicates. Numbers within the bars show the mean number of buds per replicate that entered the analysis

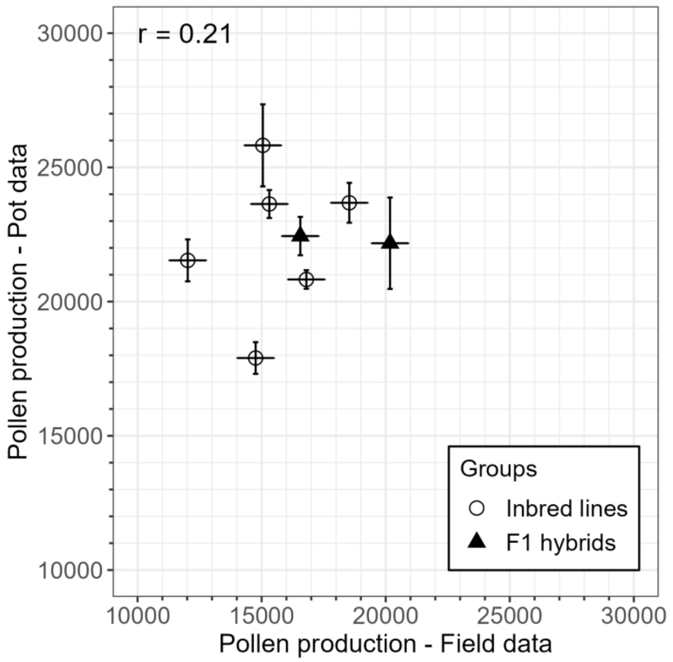

**Supplementary Fig. 7** Relationship of pollen production field data and pot data, calculated based on Spearman's product moment correlation. Error bars show standard error of the means

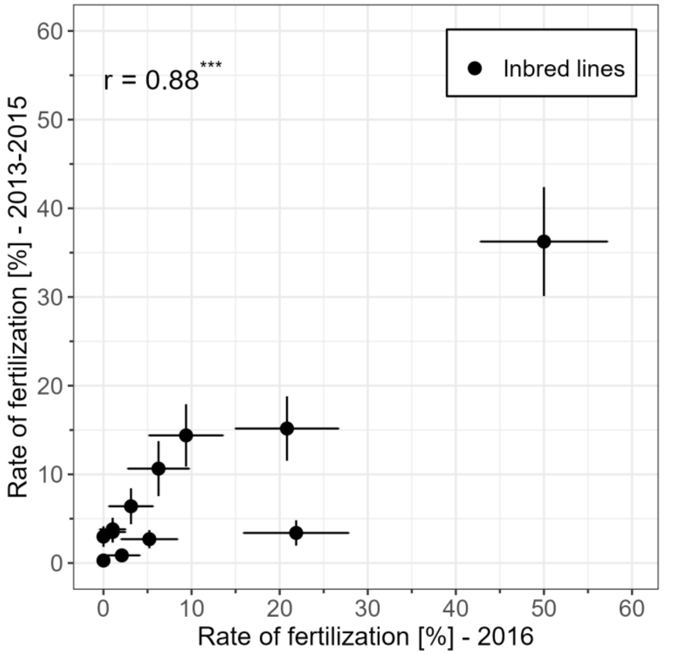

**Supplementary Fig. 8** Correlation of rate of fertilization of 2016 with the least square mean of years 2013-2015. Pearson's product moment correlation test ($t=5.93$, $df=10$, $p\text{-value} < 0.001$). Error bars show standard error of the means

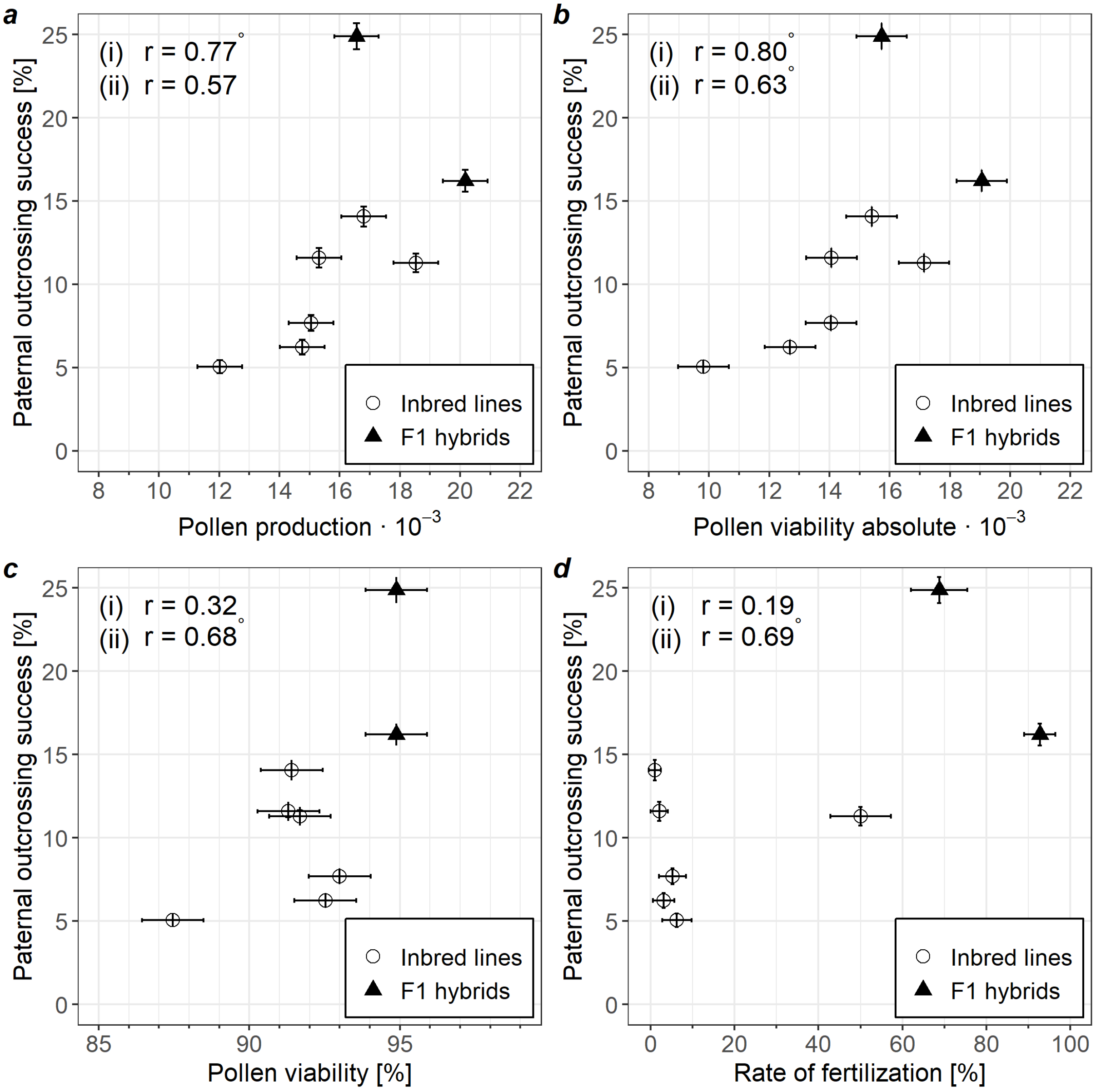

**Supplementary Fig. 9** Correlation of paternal outcrossing success with reproductive traits: **a** pollen production (field data), **b** absolute pollen viability (field data), **c** relative pollen viability (field data), and **d** rate of fertilization (pot data). Error bars show standard errors of the mean. Correlation coefficients are calculated based on Pearson's product moment correlation coefficient for (i) only inbred lines, (ii) all genotypes ( ° = significant at p = 0.1)
